# Conformational diversity of collided ribosomes determines stress signaling

**DOI:** 10.64898/2026.06.16.730951

**Authors:** Jeffrey J Li, Timo Denk, Matthias Thoms, Allen R Buskirk, Roland Beckmann, Rachel Green

## Abstract

Ribosome collisions trigger pathways that clear stalled ribosomes, and when sufficiently abundant, the integrated stress response (ISR) through GCN2 and the ribotoxic stress response (RSR) through ZAK. The inhibitors anisomycin (ANS), emetine (EME), and didemnin B (DDB) are commonly used to induce collisions in studying these responses. Here, we demonstrate that these drugs induce distinct signatures: ANS and DDB potently activate ZAK whereas EME does not. We define transcriptional programs induced by these inhibitors, where collisions induce the RSR and general inhibition of translation leads to an RSR-independent response. Surprisingly, we find that collisions induced by EME, unlike ANS, are not cleared by ASCC3. The cryo-EM structure of human disomes stalled by EME reveals its mechanism of inhibition and a conformation distinct from ANS-stalled disomes. These differences in collision geometry explain the different outcomes in quality control and signaling activation, showing how ribosome stalling events can yield distinct cellular responses.

## Introduction

Protein synthesis involves ribosomes initiating, elongating, terminating, and recycling on a messenger RNA to produce nascent proteins. However, various stresses, such as amino acid starvation, oxidative damage, or UV irradiation perturb ribosome processivity, leading to stalling and subsequent ribosome collisions^1^. Growing evidence has demonstrated that these ribosome collisions act as signals for cells to detect translational stress and activate either quality control (QC) pathways to clear the collision or stress response pathways to implement a broader cellular response^2,3^. However, it remains unclear how the cell discerns incidental collisions from those arising from translational stress.

Translational stress activates quality control pathways that degrade damaged mRNAs, rescue stalled or collided ribosomes, and target incomplete nascent polypeptides for degradation. In human cells, collided ribosomes are recognized by the E3 ubiquitin ligase ZNF598, leading to ubiquitylation of the ribosomal proteins eS10 and uS10^4–7^. These ubiquitins in turn recruit a complex containing the helicase ASCC3, which splits the lead ribosome in the collided pair into individual 40S and 60S subunits, after which the nascent polypeptide bound to the 60S subunit is degraded by the ribosome-associated quality control (RQC) pathway^8–13^.

Translational stress also activates the collision-responsive kinases GCN2 and ZAK, triggering the integrated stress response (ISR) and the ribotoxic stress response (RSR), respectively^14^. One of the four mammalian ISR kinases, GCN2 was originally identified as a sensor of amino acid starvation, which when activated, phosphorylates the translation initiation factor eIF2α, leading to a global block in translation initiation^15^. Recent work has shown that this collision-mediated activation of GCN2 limits continued collision formation during translational stress^16–18^.

ZAK is a MAP kinase kinase kinase (MAP3K) whose collision-mediated dimerization and activation occur through specific interactions of its SAM domain with the ribosomal protein RACK1 on the stalled and colliding ribosome; this disome collision structure is notably well resolved by cryo-EM of anisomycin-treated disomes suggestive of a very stable collision interface^19^. ZAK activation initiates a signaling cascade that leads to activation of the MAP kinases p38 and JNK^14,20–22^. Once activated, p38 and JNK phosphorylate their downstream targets to mediate cell cycle arrest and apoptosis, respectively, allowing cells to quickly adapt and respond to translational stress^14,16,20,23^.

To study these pathways, translation elongation inhibitors at low concentrations are frequently used to induce ribosome collisions^1,7,14,20,23,24^. Anisomycin, emetine, and didemnin B, three different elongation inhibitors with distinct inhibition mechanisms (**Figure S1A**), all induce ribosome collisions that are recognized by ZNF598^7^. However, previous studies of the RSR reported that different elongation inhibitors activate JNK and p38 with varying potencies, although the reason for these discrepancies remained unclear^20,25^. Given that the field routinely uses these elongation inhibitors to study ribosome collisions and their cellular consequences, elucidating the mechanistic basis for these differences is critical for a thorough understanding of collision-mediated signaling^7,14,20,23,26^.

Here, we demonstrate that elongation inhibitors activate QC, the ISR, and the RSR to different extents and with different dynamics even at doses that cause roughly equivalent numbers of collisions. Anisomycin, which binds the ribosomal peptidyl transferase center (PTC)^27^, and didemnin B, which traps the elongation factor eEF1A on the ribosome^28^, strongly activate the RSR. In contrast, emetine, which binds the E site of the ribosome to block translocation^29–32^, very weakly activates the RSR. We further characterize the effects of each elongation inhibitor using RNA sequencing, defining RSR-dependent and RSR-independent transcriptional programs and connecting their activation to ribosome collisions or general translation inhibition, respectively. We further show that unlike anisomycin- or didemnin B-induced collisions, emetine-induced collisions are not cleared by ASCC3. Finally, we provide a cryo-EM structure of human emetine-bound disomes, which explains their differences in QC, ISR and RSR activation relative to anisomycin treatment. Most significantly, comparison of anisomycin- and emetine-bound disomes suggests that differences in RSR signaling are driven by structural differences in collision conformation, wherein the RACK1-RACK1 spacing on the emetine disome is greater and substantially looser than that observed in the anisomycin disome, thus disfavoring ZAK activation by this collision interface. More broadly, these findings highlight the fact that not all stalled ribosomes are quickly cleared by rescue factors and not all collisions trigger stress signaling pathways.

## Results

### Different elongation inhibitors exhibit distinct effects on RSR activation

Treatment with sub-stoichiometric doses of elongation inhibitors is a well-established method to induce ribosome collisions^1^. Given that inhibitors target different stages of the translation elongation cycle (**Figure S1A**), we wondered whether ribosome collisions induced by these inhibitors behave similarly or show differences in their lifetimes or in their effects on downstream signaling pathways. We first titrated the concentration of anisomycin (ANS), emetine (EME), and didemnin B (DDB) in HEK293T cells and visualized the levels of ribosomal protein eS10 ubiquitylation, a classical marker for ribosome collisions^4–6^ after 15 min of treatment (**Figure S1B**). As expected, eS10 ubiquitylation exhibited dose dependence in each case, with maximal ubiquitylation occurring at intermediate doses of each drug, as previously reported^1^. We adopted these doses (0.38 μM ANS, 1.8 μM EME, and 0.5 μM DDB) as our “standard” collision-inducing concentrations for the remainder of the study. At these doses, ANS and EME induce eS10 ubiquitylation to a similar degree, while DDB induces lower amounts (**Figure S1B, bold**). Nonetheless, all three inhibitors trigger eS10 ubiquitylation above basal levels, suggesting that these drugs induce ribosome collisions that recruit ZNF598 despite their different modes of translational inhibition^7^.

Another hallmark of collisions is that nucleases, such as RNase A, cannot cleave the mRNA between collided ribosomes. To compare the number of collisions generated by each condition, we digested drug-treated lysates with RNase prior to sucrose gradient fractionation. As expected, treatment with any of the three inhibitors for 15 min led to comparable increases in the amount of nuclease-resistant disomes (**Figure S1C, top**). These data are consistent with the idea that these chosen doses of the three inhibitors induce roughly similar levels of ribosome collisions at 15 min, allowing us to compare their effects on downstream signaling pathways.

Ribosome collisions trigger ZAK autophosphorylation, subsequent dissociation from the ribosome, and phosphorylation of substrates p38 and JNK in the ribotoxic stress response^14,16,19^. To evaluate the ability of each inhibitor to activate the RSR, we treated HEK293T cells with ANS, EME, or DDB for either 15 min or 2 h prior to immunoblotting for signatures of RSR activation (**Figure 1A**). We utilized a Phos-tag SDS-PAGE approach to observe ZAK activation directly: phosphorylated ZAK migrates more slowly and appears as a smear above unphosphorylated protein^33^. In line with previous studies^14,20^, we observed ZAK phosphorylation with ANS or DDB treatment, but not EME (**Figure 1A**), despite comparable levels of eS10 ubiquitylation and nuclease-resistant disomes (**Figure S1C**) across these conditions. To further define inhibitor-specific differences in ZAK activation, we looked at phosphorylation of JNK and p38, the downstream targets of ZAK in the RSR cascade; again, we observed strong induction of JNK and p38 phosphorylation with ANS and DDB, while little or no JNK or p38 phosphorylation was observed with EME (**Figure 1A**). These observations are consistent with early studies of the RSR reporting that ANS induces a stronger RSR response than EME^25^. Interestingly, while ANS and DDB both strongly activated the RSR, these two inhibitors exhibited differences in the timing of RSR activation. ANS potently induced JNK and p38 activation at 15 min, and this signal decayed significantly after 2 h; in contrast, DDB weakly induced JNK and p38 activation by 15 min but activation increased to higher levels at 2 h. Taken together, these data suggest that all three inhibitors exhibit distinct patterns of RSR activation: (1) ANS induces strong but transient RSR activation; (2) EME induces barely detectable RSR activation; (3) DDB gradually induces sustained RSR activation. Importantly, for all three inhibitors, drug-induced phosphorylation of JNK and p38 was strongly blocked by pre-treatment with nilotinib, a ZAK inhibitor (ZAKi) (**Figure S1D**), confirming that these phosphorylation events are primarily driven through the RSR. Taken together, our data demonstrate that robust formation of ribosome collisions is not the sole determinant of RSR activation.

**Figure 1:**
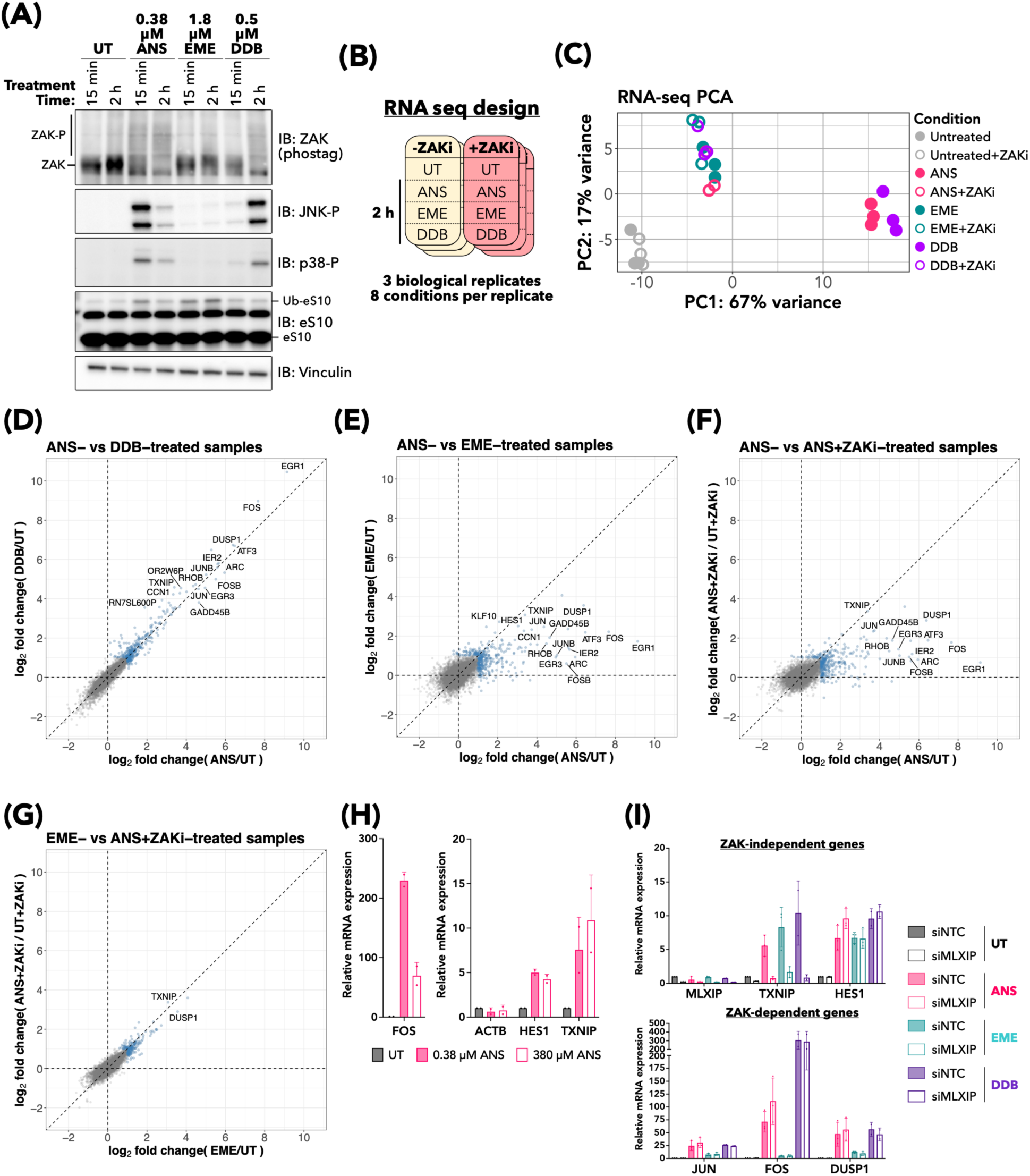
Ribosome collisions induced by different elongation inhibitors exhibit distinct patterns of ribotoxic stress response activation. **(A)** Immunoblots of HEK293T cells left untreated (UT) or with the indicated dose and times of anisomycin (ANS), emetine (EME), or didemnin B (DDB). **(B)** Schematic of RNA-sequencing experiment design. HEK293T cells were pre-treated with DMSO or ZAK inhibitor for 1 h prior to treatment with elongation inhibitor for 2 h. **(C)** Principal component analysis (PCA) plot of RNA-seq samples. Samples are annotated both by color (indicating elongation inhibitor treatment) and fill (indicating ZAK inhibitor pre-treatment). **(D-G)** Cross-correlation plots of gene expression in the indicated inhibitor-treatment conditions normalized to the indicated untreated condition. Points highlighted in blue indicate genes showing a log_2_(fold change) (log_2_FC) greater than 1 in at least one condition. **(H)** RT-qPCR of HEK293T cells left untreated or treated with 0.38 µM or 380 µM ANS for 2 h.**(I)** RT-qPCR of HEK293T cells transfected with non-targeting control (siNTC) or MLXIP-targeting (siMLXIP) siRNAs prior to treatment with 0.38 µM ANS, 1.8 µM EME, or 0.5 µM DDB for 2 h.

### Defining the ZAK-dependent and -independent transcriptional responses via RNA-seq

We next asked how inhibitor-specific differences in RSR activation affect transcriptional responses. To this end, we performed RNA-seq on HEK293T cells pre-treated with DMSO or ZAK inhibitor followed by 2 h treatment with ANS, EME, or DDB (**Figure 1B**). Principal component analysis (PCA) demonstrated clear separation of the samples into discrete clusters according to their treatment condition (**Figure 1C**). ANS- and DDB-treated samples clustered together and strongly separated away from both the untreated and EME-treated sample clusters in line with inhibitor-specific differences in RSR activation. In contrast, the EME-treated samples, independent of pre-treatment with ZAK inhibitor, occupied a unique cluster distinct from both the untreated cluster and the ANS-/DDB-treated cluster. Given that EME barely activates the RSR (**Figure 1A**), these data suggest that EME treatment induces a distinct transcriptional response that is independent of the ZAK-mediated RSR. Interestingly, when pre-treated with ZAK inhibitor, the ANS and DDB samples (ANS+ZAKi and DDB+ZAKi) grouped near the EME cluster, rather than near the untreated cluster, suggesting that ANS and DDB also trigger an RSR-independent transcriptional program. Thus, our RNA-seq dataset captures distinct ZAK-dependent and ZAK-independent contributions to inhibitor-induced transcriptional changes.

We next explored these trends and identified which genes are shared between each inhibitor’s transcriptional program. The gene expression changes observed upon ANS and DDB treatment were highly correlated, suggesting these two drugs activate the same transcriptional program (**Figure 1D**). In contrast, comparing the ANS or DDB samples with EME revealed a large subset of genes that were highly induced only by ANS or DDB (**Figures 1E, S2A**); we hypothesized that these genes represent the RSR transcriptional program and thus should be ZAK-dependent. Consistent with this hypothesis, ANS-induced or DDB-induced expression of these genes is strongly inhibited by pre-treatment with ZAK inhibitor (**Figures 1F, S2B**). Strikingly, the ANS sample pre-treated with ZAK inhibitor closely resembles the EME treated sample (**Figure 1G**), suggesting that genes induced under these conditions represent a ZAK-independent transcriptional program activated in response to translational stress.

Detailed analysis of the RNA-seq dataset with DESeq2 revealed that 296 and 286 genes were significantly up-regulated (> 2-fold & p_adj_ < 0.01) by ANS and DDB respectively (**Figure S2F**). These induced genes exhibit strong overlap between the two conditions, consistent with both inhibitors inducing similar transcriptional programs (**Figure S2F, upper**). This ANS- and DDB-mediated program decreased to 111 and 126 genes upon ZAK inhibitor pre-treatment (**Figure S2F, lower**). In contrast, the number of EME-induced genes did not significantly change with ZAK inhibitor pre-treatment, with 153 and 161 genes up-regulated in the absence and presence of ZAK inhibitor respectively (**Figure S2F**). Following ANS or DDB treatment, the ZAK-dependent genes, such as *EGR1* and *FOS,* show the highest fold changes (**Figures 1F, S2B**), highlighting the importance of ZAK in the transcriptional response downstream of collision stress. Notably, many of these highly expressed ZAK-dependent genes such as *JUN*, *FOS*, and *ATF3* were also identified in early studies describing the induction of immediate-early genes following ribotoxic stress^25,34,35^. In contrast, none of the EME-induced genes achieved similar fold changes in expression as those induced by ANS or DDB (**Figures 1E, S2A**). For instance, the highest expressing gene induced by ANS is *EGR1*, exhibiting a 561-fold increase in expression compared to untreated conditions (**Figure S3A**); conversely, the highest expressing gene induced by EME, encoding a putative Histone H2A isoform (ENSG00000282988), exhibits only a 17-fold increase in expression (**Figure S3E**). Interestingly, comparison of the ZAK inhibitor treatment conditions showed significant overlap across all three drugs, suggesting that all three inhibitors trigger a similar ZAK-independent transcriptional program (**Figure S2F, lower**).

To further characterize the transcriptional responses induced by each inhibitor, we conducted gene set enrichment analysis (GSEA) of our RNA-seq datasets. In all three conditions, genes involved in broad signaling gene sets such as “TNF-α signaling via NF-κB”, “apoptosis”, and “hypoxia” were enriched (**Figures S4A-C, solid dots**). This enrichment was also observed even with pre-treatment with the ZAK inhibitor, suggesting a broad ZAK-independent response to translation inhibition (**Figures S4A-C, hollow dots**). However, enrichment of the “UV response up” gene set was blocked by the ZAK inhibitor in the ANS and DDB conditions (**Figure S4A, S4C**), consistent with our group’s previous report defining the importance of ZAK in the UV-mediated RSR^16^. Importantly, in line with our PCA, GSEA of the ANS and DDB conditions showed remarkable concordance in gene sets enriched (**Figure S4D**), whereas the EME condition alone showed de-enrichment of gene sets involved in metabolism, such as “glycolysis” and “oxidative phosphorylation” (**Figure S4B, S4D**). Finally, in the presence of ZAK inhibitor, ANS, EME, and DDB all exhibited similar patterns of enriched and de-enriched gene sets (**Figure S4E**), suggesting that absent ZAK activation, all three inhibitors induce similar transcriptional programs.

### ZAK-independent transcriptional changes are driven by general translation inhibition

Since ANS, EME, and DDB all exhibit similar gene expression changes when ZAK is inhibited (**Figures 1C, S2F**), we hypothesized that these changes represent ZAK-independent signaling driven by general translation inhibition, rather than ribosome collisions. To evaluate this hypothesis, we conducted RT-qPCR for RSR and non-RSR transcripts on cells treated with either 0.38 µM or 380 µM ANS, which lead to widespread ribosome collision or stalling, respectively (**Figure 1H**). However, as shown in previous reports^20^, we note that high doses of ANS do not completely prevent collisions (**Figure S1B**). Nevertheless, *FOS*, a gene strongly induced by the RSR, demonstrates dose-dependent induction (**Figure 1H - left**), while *TXNIP* and *HES1*, two ZAK-independent transcripts identified in our RNA-seq dataset, demonstrate dose-independent induction (**Figure 1H - right**). These data indicate that induction of these ZAK-independent genes does not correlate with collision number and is likely due to general inhibition of translation elongation rather than ribosome collisions per se.

We initially chose to use a ZAK inhibitor with high potency and specificity^16^ rather than a genetic knock-out to avoid potential issues associated with clonal variability and genetic compensation. However, we wondered to what degree observed transcriptional changes could be driven by residual ZAK activity despite pre-treatment with the ZAK inhibitor. To test this, we treated wildtype (WT) and monoclonal ZAK-deletion (ΔZAK) HEK293T cell lines with ANS or DDB, which potently activate ZAK-mediated transcription, and monitored the expression of ZAK-dependent and ZAK-independent targets by RT-qPCR. Expression of the stress response genes *JUN*, *FOS*, and *DUSP1* upon ANS or DDB treatment was nearly abolished in the ΔZAK cell line (**Figure S5A**). Thus, it is likely residual ZAK activity, rather than a ZAK-independent mechanism, that explains the induction of these particular genes with ANS or DDB in the presence of ZAK inhibitor. In contrast, *HES1* and *ARRDC3*, two ZAK-independent genes identified in our RNA-seq experiment (**Figures S3B, S3D, S3F**), did not show a difference in drug-mediated up-regulation between the WT and ΔZAK cell line (**Figure S5A**), suggesting that their induction is indeed ZAK-independent. Interestingly, *TXNIP* up-regulation is reduced in the ΔZAK cell line, suggesting that its induction depends at least in part on ZAK activity (**Figure S5A**).

We next asked what drives the expression of ZAK-independent genes downstream of translation inhibition by these diverse agents. Our RNA-seq study demonstrated that *TXNIP* is one of the most up-regulated transcripts in samples pre-treated with ZAK inhibitor (**Figures S3B, S3D, S3F**). Known as a negative regulator of glucose uptake, TXNIP has been shown to be strongly induced by translation inhibitors through the activity of the transcription factor MLXIP that senses changes in cellular ATP^36,37^. Thus, we hypothesized that the action of MLXIP may explain part of the ZAK-independent signaling downstream of elongation inhibition; indeed, knockdown of MLXIP potently blocked transcriptional induction of *TXNIP* upon treatment with ANS, EME, or DDB (**Figure 1I**). In contrast, transcription of the ZAK-dependent stress response genes *JUN, FOS*, and *DUSP1* was completely unaffected by MLXIP knockdown. Interestingly, transcription of *HES1*, a gene significantly up-regulated in a ZAK-independent manner, was also unaffected by MLXIP knockdown. Overall, these data suggest that the gene expression program activated downstream of general inhibition of protein synthesis is not driven by a single transcription factor.

### EME blocks clearing of ribosome collisions by ASCC3

Based on our observations that ANS, EME, and DDB treatment all lead to ribosome collisions but activate the RSR to different extents, we asked what might explain these differences. We began by investigating the effects of each inhibitor on the translational state of the cell using polysome profiling, treating cells with collision-inducing doses of each drug for either 15 min or 2 h. ANS treatment led to a striking time-dependent accumulation of 80S monosomes and free 60S subunits, with a corresponding decrease in polysomes over time (**Figure 2A, left**). In contrast, EME treatment led to a minor decrease in 80S monosomes and a time-dependent increase in heavy polysomes (**Figure 2A, middle**). Finally, DDB treatment led to an 80S monosome accumulation, similar to that observed with ANS, though this DDB-induced 80S increase was only seen at the 2 h timepoint and was not associated with an increase in 60S subunits (**Figure 2A, right**). Additionally, with 2 h DDB treatment, there was a shift in polysomes from predominantly heavy fractions towards lighter fractions. Thus, the three inhibitors have distinct effects on translation.

**Figure 2:**
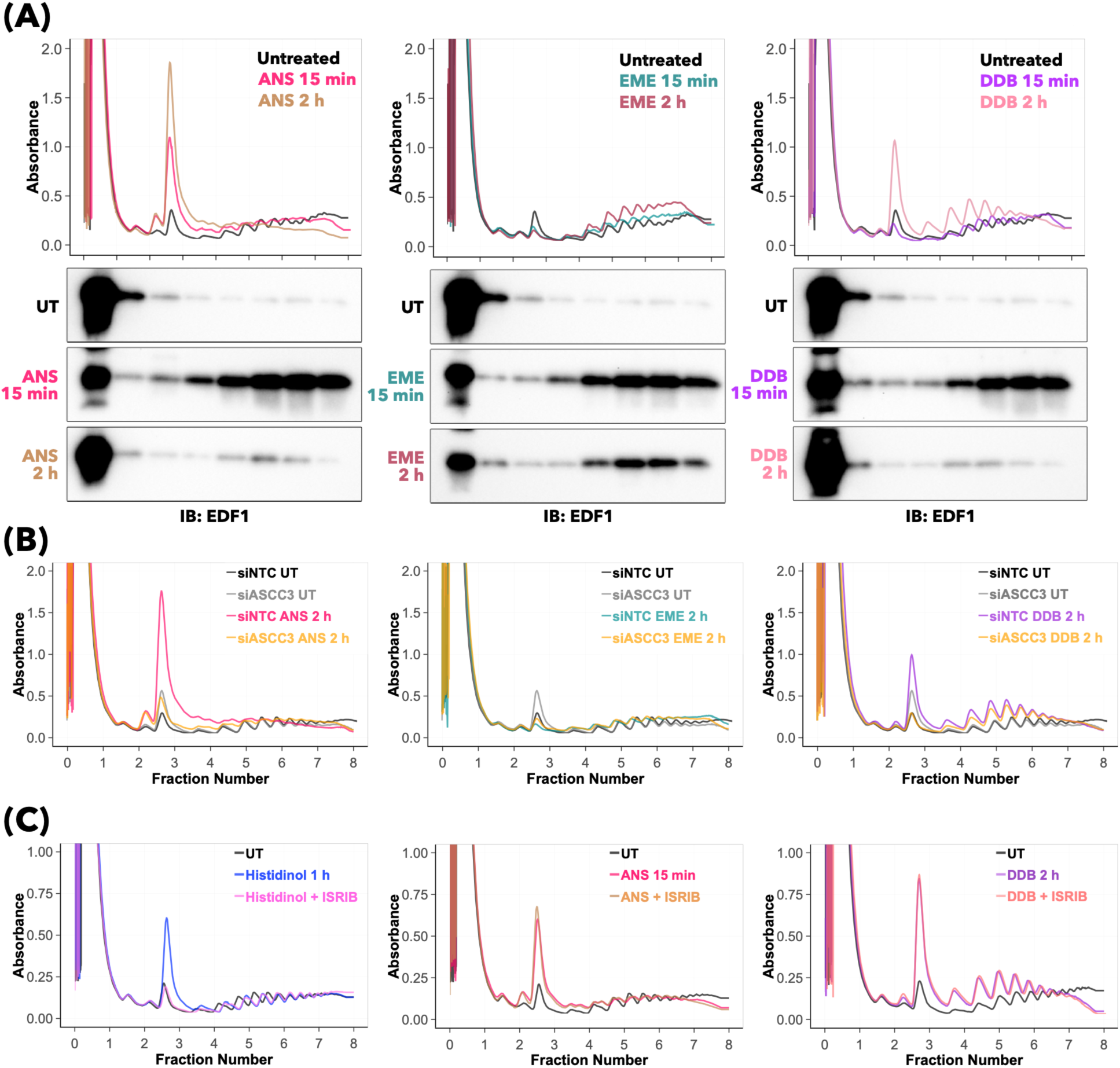
Different elongation inhibitors exert distinct effects on translation reinitiation and collision clearance. **(A)** Polysome traces of HEK293T cells left untreated or treated with anisomycin (ANS), emetine (EME), or didemnin B (DDB) for 15 min and 2 h. Fractions were immunoblotted for EDF1. The same polysome trace and immunoblot of the untreated condition is shown across all three condition sub-panels for clarity. **(B)** Polysome traces of HEK293T cells transfected with non-targeting (siNTC) or ASCC3-targeting (siASCC3) siRNAs prior to treatment with 0.38 µM ANS, 1.8 µM EME, or 0.5 µM DDB for 2 h. The same polysome traces of the untreated conditions are shown across all three condition sub-panels for clarity. **(C)** Polysome traces of HEK293T cells left untreated or pre-treated with ISRIB for 1 h prior to treatment with 4 mM histidinol for 1 h, 0.38 µM ANS for 15 min, or 0.5 µM DDB for 2 h. The left and middle panels originated from the same experiment, and thus the corresponding untreated polysome trace is the same for the left and middle panels.

We used EDF1 recruitment into polysomes as a proxy^24,38^ for collision accumulation. Treatment with any of the three inhibitors for 15 min led to substantial EDF1 recruitment into the polysome fractions, indicative of pervasive collisions (**Figure 2A**). After 2 h of ANS or DDB treatment, EDF1 returned to the free fraction, suggesting that ribosome collisions are largely resolved by this timepoint (**Figure 2A, left/right**); in contrast, a substantial amount of EDF1 remained bound to polysomes even with prolonged EME treatment (**Figure 2A, middle**). These data suggest either that EME, but not ANS or DDB, induces collisions that are resistant to clearance machinery, or alternatively that EME uniquely allows ongoing translation and continual collision formation at extended timepoints. Overall, these observations are consistent with our earlier data on eS10 ubiquitylation (**Figure 1A**): ANS potently induces eS10 ubiquitylation within 15 min that decays by 2 h, consistent with the shift of polysome-bound EDF1 back into the free fraction. In contrast, EME induces similar levels of eS10 ubiquitylation at 15 min that persist after 2 h. This persistence is also consistent with our nuclease-resistant disome data, where unlike ANS or DDB, EME induces similar levels of disomes at 15 min that remain after 2 h treatment (**Figure S1C, bottom**). These observations suggest that ANS-induced and DDB-induced collisions are cleared more quickly than EME-induced collisions.

A previous study also observed an ANS-induced 80S accumulation and showed that it was ZNF598 dependent^26^. Given ZNF598’s importance in sensing collisions and triggering quality control pathways, we next tested the idea that the clearing of collided ribosomes by ASCC3 is responsible for the decrease in polysomes and the corresponding increase in 80S monosomes induced by ANS and DDB treatment. Knockdown of ASCC3 with siRNAs led to a dramatic reduction in 80S accumulation following treatment with ANS (**Figure 2B, left**) or DDB (**Figure 2B, right**) suggesting that the 80S accumulation is indeed due to clearing of collided ribosomes. In contrast, regardless of whether ASCC3 is knocked down or not, EME treatment does not lead to 80S accumulation (**Figure 2B, center**). Notably, ASCC3 knockdown leads to RSR and ISR hyperactivation even without the addition of elongation inhibitors (**Figure S6A**), suggesting that collision clearance generally negatively regulates RSR and ISR activation as previously reported^17,18,23^.

While the loss of polysome-bound EDF1 and the ASCC3-dependent accumulation of 80S ribosomes argues that collisions are cleared in the ANS and DDB samples, the EME results are more challenging to interpret. On the one hand, it may be that EME-induced collisions are resistant to clearance by ASCC3, but it is also possible that ribosomal subunits from cleared collisions rapidly reinitiate to form new collisions in a dynamic cycle. To distinguish between these possibilities, we treated cells with EME for 15 min to induce collisions before acutely blocking ongoing ribosome loading with sodium arsenite, which induces a potent initiation block through activation of the integrated stress response (ISR). Under these conditions, if EME-induced collisions were being cleared by ASCC3, the resulting ribosomal subunits should not be able to re-initiate and thus should accumulate in the 80S peak as seen with ANS treatment. As controls, we observed that EME treatment alone led to an accumulation of heavy polysomes (**Figure S6B, green**), while arsenite treatment alone led to a complete loss of polysomes and subsequent 80S accumulation (**Figure S6B, blue**). Importantly, the EME-arsenite chase experiment did not result in the accumulation of 80S monosomes or a loss of polysomes (**Figure S6B, pink**), strongly suggesting that EME-induced collisions are resistant to clearance by ASCC3.

### Elongation inhibitor treatment does not block initiation through the ISR

We next wondered why the ribosomes cleared by ASCC3 in the ANS and DDB samples do not re-initiate and re-enter the translational cycle (**Figure 2A**). This phenotype, with the large 80S peak, is reminiscent of ISR activation^39^ and would be consistent with previous work suggesting that ANS induces the ISR to block translation initiation by activating the kinase GCN2^14^. To evaluate this possibility, we pre-treated cells with ISRIB, a well-established ISR inhibitor^40^, prior to treatment with histidinol or ANS. Histidinol inhibits histidine tRNA charging, leading to a histidine starvation phenotype^41^ and potent activation of the ISR through GCN2. As expected, histidinol induced an 80S accumulation characteristic of the ISR^39^, and this effect was completely blocked by ISRIB pre-treatment (**Figure 2C, left**); in contrast, ANS or DDB induced 80S accumulation but was unaffected by ISRIB pre-treatment (**Figure 2C, center and right**). These data indicate that ANS or DDB do not potently activate the ISR through GCN2. We also considered the possibility that 80S accumulation is dependent on RSR activation, since the timing of 80S accumulation correlated with that of ZAK phosphorylation (**Figures 1A, 2A**). However, pre-treatment with ZAK inhibitor had no effect on ANS-induced 80S accumulation (**Figure S6C**). We conclude that the observed ANS-induced accumulation of 80S ribosomes is downstream of ASCC3-mediated collision clearance, but is not the result of activation of the ISR or the RSR.

### Inhibitor-specific differences in RSR activation are not driven by differences in collision number

We initially aimed to identify ANS, EME, and DDB doses that induce similar levels of collisions, using eS10 ubiquitylation (**Figure S1B**) and nuclease-resistant disome profiling (**Figure S1C**) as collision abundance metrics. In addition, we note that at our chosen doses (0.38 µM ANS, 1.8 µM EME, 0.5 µM DDB), 15 min of treatment with any of the three drugs led to similar amounts of EDF1 recruitment to polysomes (**Figure 2A**). However, based on the differences in clearing by ASCC3 described above, we wondered if the differences in RSR activation were driven by undetected differences in collision number integrated over time. It is possible that ANS- or DDB-induced collisions activate the RSR by enabling ZAK activation at the collision interface, but the collisions are quickly cleared by ASCC3, such that the overall number of collisions experienced during a time period is not reflected in the “steady state” number of collisions observed at a single timepoint as measured by eS10 ubiquitylation or RNase resistant disome levels. In other words, the RSR activation we observe may be the consequence of the accumulation of collision events during treatment, rather than the number of collision events observed at a given time. Additionally, the eS10 ubiquitylation and RNase resistant disome assays have inherent shortcomings, as eS10 ubiquitylation may partially correspond to already cleared 40S subunits awaiting deubiquitylation^42^ while nuclease-resistant polysome profiling is prone to over-digestion and therefore can underestimate collision abundance.

To address these limitations, we developed an EDF1 recruitment assay that allowed us to more directly measure collision levels with higher sensitivity. EDF1 specifically binds collided ribosomes^24,38^ and only enters the ribosome-containing pellet fraction when ribosome collisions are present. Using our optimized conditions defined in Figure 1, 15 min treatment with any of the three inhibitors at “standard” collision-inducing doses leads to a nearly complete shift of EDF1 from the supernatant to the ribosome-containing pellet (**Figure 3A**). These data suggest that all three drugs potently induce ribosome collisions at the 15 min timepoint.

**Figure 3:**
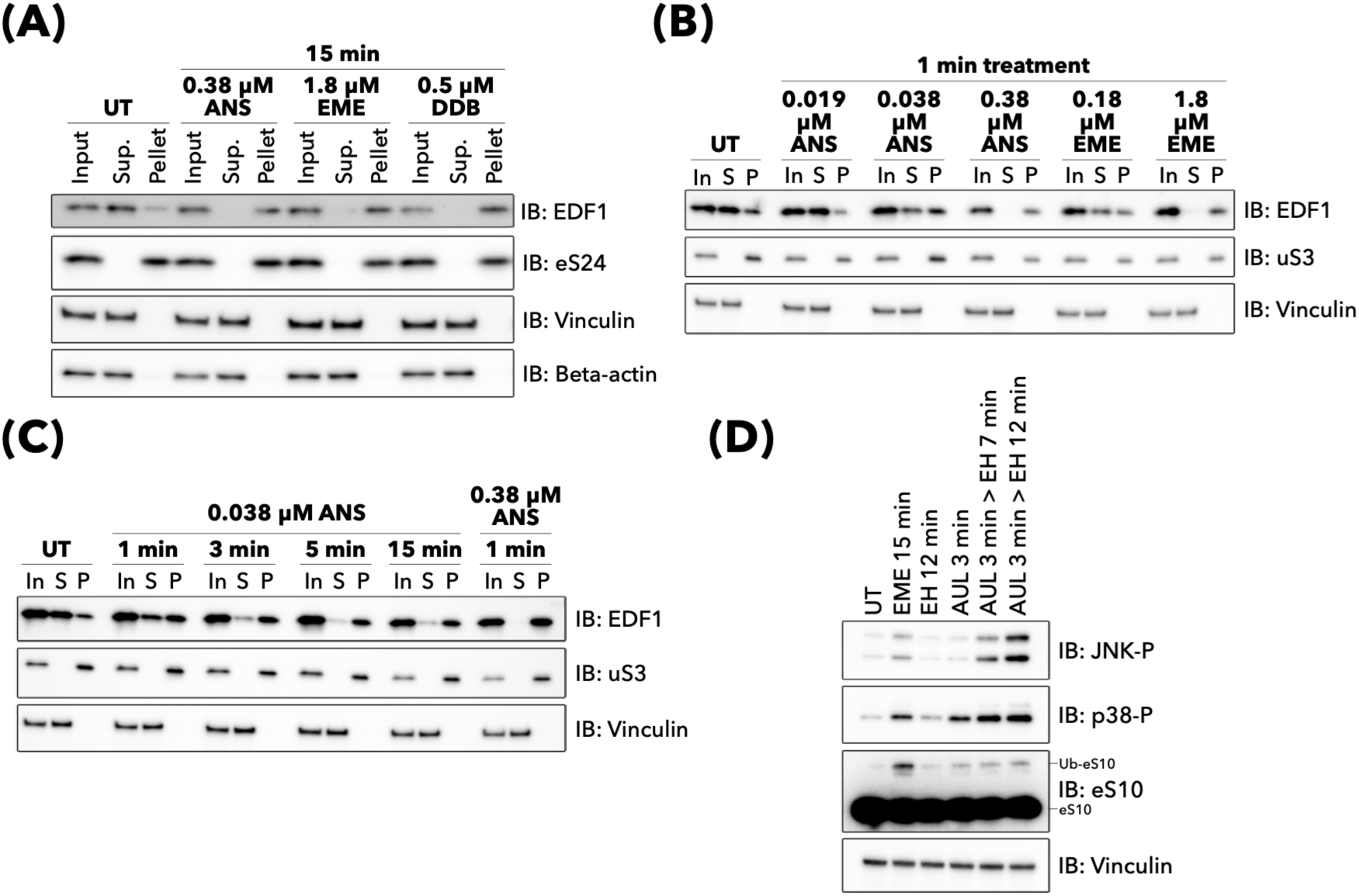
Inhibitor-specific differences in RSR activation are not due to differences in collision number. **(A)** Immunoblots of the input, supernatant (Sup), and pellet fractions after ribosome pelleting of HEK293T cells left untreated (UT) or with the indicated dose of anisomycin (ANS), emetine (EME), or didemnin B (DDB) for 15 min. **(B)** Immunoblots of the input (In), supernatant (S), and pellet (P) fractions after ribosome pelleting of HEK293T cells left untreated (UT) or with the indicated dose of ANS or EME for 1 min. **(C)** Immunoblots of the input (In), supernatant (S), and pellet (P) fractions after ribosome pelleting of HEK293T cells left untreated (UT) or with the indicated dose of ANS for the indicated times. **(D)** Immunoblots of HEK293T cells left untreated (UT), treated with 1.8 µM EME (EME) for 15 min, 90 µM EME (EH) for 12 min, 0.038 µM ANS (AUL) for 1 min, or 0.038 µM ANS (AUL) for 3 min prior to addition of 90 µM EME (> EH) for 7 min or 12 min to freeze translation.

However, cellular proteome measurements estimate that ribosomes outnumber EDF1 by about 5:1^43^. Thus, complete EDF1 recruitment to the ribosome pellet cannot report on precise collision abundance. To address this, we titrated ANS and EME to a more sensitive range where EDF1 is not completely recruited to the ribosome pellet, allowing us to more accurately compare collision levels in different regimes. With 1 min of treatment, decreasing our standard dose of ANS or EME by 10-fold led to a partial, rather than complete, shift of EDF1 into the ribosome pellet (**Figure 3B**). A treatment time course of this “ultra-low” dose of 0.038 µM ANS (AUL) revealed that EDF1 shifts from the supernatant to the pellet fraction over time (**Figure 3C**), consistent with the idea that collisions accumulate during treatment. Importantly, at all times tested for the “ultra-low” ANS dose, a fraction of EDF1 remained in the supernatant (**Figure 3C**). This demonstrates that these “ultra-low” doses induce fewer collisions than our standard dose of inhibitor, which fully recruits EDF1 to ribosomes even after 1 min of treatment.

We next sought to understand how rapidly collision clearance and RSR activation occur by treating cells with shorter drug incubations (< 15 min). Polysome profiling showed that ANS-mediated 80S accumulation, which is ASCC3-dependent (**Figure 2B**), is only observable after 5 min (**Figure S7A**), indicating a delay between collision formation and ASCC3 activity. However, ANS-induced eS10 ubiquitylation reached near-maximal levels after 1 min treatment (**Figure S7B**), suggesting that ZNF598 is able to rapidly recognize collisions. RSR activation is comparatively slower, as JNK and p38 activation was weakly observed at 5 min and only reached maximal potency at 10 min treatment (**Figure S7B**). Overall, these data suggest that after collision formation, there is a delay before both ASCC3-mediated collision clearance and ZAK-mediated RSR activation can be observed.

With these parameters in place, we asked if a sub-maximal amount of ANS-induced collisions, as measured by incomplete recruitment of EDF1 to ribosomes, can still trigger the RSR (**Figure 3D**), as compared with the higher “optimal collision inducing” dose of EME. Since we hypothesized that drug-induced collision accumulation may differ over time due to differences in ASCC3 activity, we chose short treatments (< 3 min) to minimize the effect of ASCC3-mediated collision clearance (**Figure S7A**). Additionally, since the RSR signal takes time to develop (**Figure S7B**), we followed a short treatment of “ultra low” dose ANS (AUL) with a high dose of EME (EH) to inhibit further elongation and thus further collision formation while giving time for ZAK activated during the initial AUL treatment to propagate its signaling cascade. Consistent with our previous findings, 1.8 µM EME, which triggers complete EDF1 recruitment to ribosomes (**Figure 3B**), induced very low levels of JNK and p38 activation even after 15 min of treatment; in contrast, 3 min of “ultra-low” dose ANS treatment (AUL) produced potent JNK activation that developed by the same 15 minute time point (**Figure 3D**). This AUL-driven effect occurs despite less eS10 ubiquitylation compared to EME (**Figure 3D**) as well as incomplete EDF1 recruitment to ribosomes (**Figure 3C**). Overall, these data support the idea that the inhibitor-specific differences in RSR signaling that we observe are not driven by differences in collision number.

### EME treatment induces the formation of non-canonical disomes

To understand what causes the distinct stress response activation between EME and ANS/DDB treatment, we chose a structural approach to investigate potential molecular differences in ribosome collisions formed by EME. We treated human FreeStyle 293-F cells with high dose EME (100 µM) for 15 min to ensure robust drug binding (but anticipating decreased collision abundance) and directly applied cell lysate to cryo-EM grids. This *in extracto* cryo-EM method bypasses lengthy isolation procedures and allows for near-unperturbed preservation of *in situ* translation state distributions^44^. At the same time, it grants the high-resolution advantages of single-particle processing compared to cryo-electron tomography (cryo-ET). Initial 3D classification showed that the distribution of ribosomal states after EME treatment was skewed towards a majority of particles with eEF2 bound and additional density at the mRNA exit indicative of a trailing ribosome (**Figure S8**). This also was the sole class of ribosomes that contained EME. All other ribosomal classes lacking EME presented with additional weak density at the mRNA entry, hinting towards the presence of a subpopulation of particles containing leading ribosomes. By an iterative series of box expansion and classification steps using the EME-bound ribosomal particles, it became clear that the collision arrangement was not rigid but rather dynamic (**Figure S8C**). By applying the same processing procedure (box expansion, etc.) to the class of collided, hybrid PRE-translocation ribosomes, we were able to jointly reconstruct a low-resolution disome. This, in turn, allowed us to backfit the initial maps for the EME-stalled and - collided ribosome classes prior to extensive sorting, and to subsequently obtain a high-resolution composite disome reconstruction and model (**Figure S8C and S9**).

In this disome arrangement, we found the EME-stalled 80S bound by eEF2 and locked in the final steps of translocation (TI-4/TI-5) (**Figures 4A-D**). Here, the tRNAs were in ap2/P and pe2/E transition states as observed before in cryo-EM structures of yeast translocation intermediates (**Figure 4C,D**)^45^. The collided ribosome was in a classic hybrid pre-translocation state with A/P P/E tRNAs. As expected and consistent with our data and previous studies^24^ on EME-induced collisions, EDF1 was associated primarily with the collided ribosome (**Figure 4C**).

**Figure 4:**
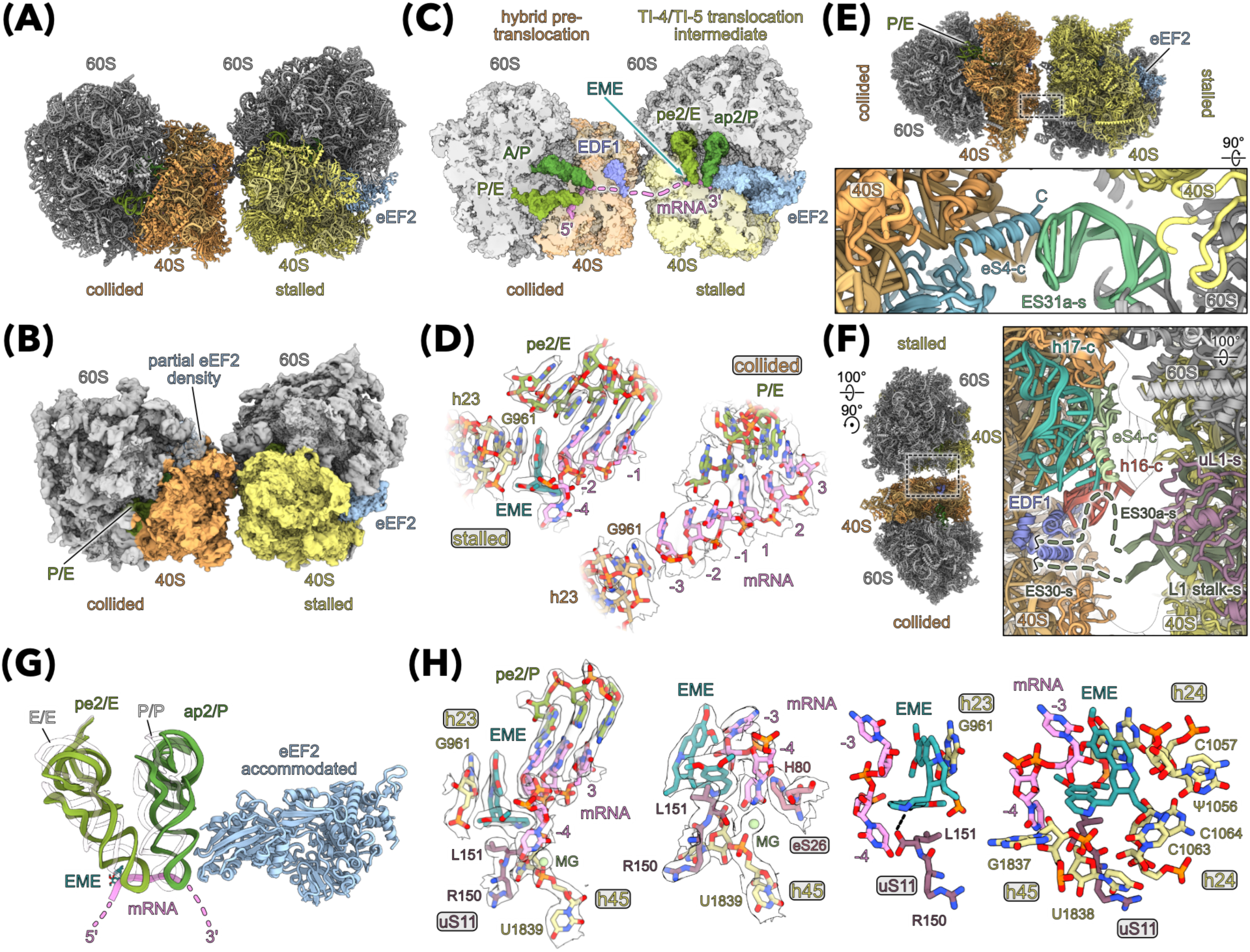
Structure of the EME disome and details of EME binding. **(A and B)** Molecular model (A) and cryo-EM density (B) of the EME disome. **(C)** Cut-through view of the EME disome focusing on tRNA states shown as surface representations. **(D)** Focused view on the mRNA and E site in stalled and collided 80S with zoned density. **(E and F)** Contact sites of the EME disomes. Focused views of the 60S-s (stalled ribosome) and 40S-c (collided ribosome) (E) and the L1-stalk-s and 40S-c (F) contacts. **(G)** Comparison of tRNAs in the EME stalled translocation intermediate with the classic POST state (PDB:9P72). **(H)** Details of the EME binding site and interactions with mRNA, ribosomal proteins and rRNA. Molecular models and zoned densities are shown.

Interestingly, the combination of late translocation intermediate stalled and hybrid collided ribosomes results in a unique disome geometry. Here, the two 80S ribosomes are further separated (**Figure 4A-C**) compared to canonical stable collisions which consist of a stalled POST and a collided hybrid state ribosome, as previously observed in mammalian cells downstream of different stalling triggers^7,19,46^. In line with this observation, the canonically tight collision interface is instead limited to two modest contact sites for the EME disome. One site consists of the 28S rRNA expansion segment ES31a-s (from the stalled (-s) 60S) that contacts the 40S ribosomal protein eS4-c (**Figure 4E**); interestingly, the interaction between ES31a-s and eS4-c is part of the canonical disome interface^7,46^. However, in the canonical disome, ES31a-s is slightly displaced and makes additional contacts with the 18S expansion segment ES6a-c. The second contact site is only observed at lower contour levels and is mediated by the L1-stalk expansion segment ES30a-s of the stalled ribosome (**Figure 4F**) which appears to contact the area around the 18S rRNA helices h17-c and h16-c as well as the N-terminal α-helix of eS4-c of the collided ribosome. This particular L1-stalk interaction is also visible in collisions observed by *in situ* cryo-ET and might be considered a more common interaction^26^.

When looking at the molecular cause for the EME disome arrangement, it becomes evident that the binding of EME into the E site on the 40S body displaces the mRNA from its usual interaction site at G961 of the 18S rRNA helix h23 (**Figure 4D**). This blocks the transition of the tRNAs from A to P, and P to E sites, which is mediated by translation elongation factor eEF2 (**Figure 4G**). In addition, EME appears to lock the mRNA in place by the formation of an extensive interaction network with several ribosomal proteins and rRNA moieties participating: by stacking in between h23 G961 and the -3 mRNA base, EME forces the -4 base into a pocket (**Figure 4H left**). Here, the -4 base is held in place by interaction with eS26, EME, and an interaction with a magnesium coordinated by the h45 U1839 phosphate. EME itself is additionally fixed by interactions with the C-terminus of uS11 and the 18S rRNA helices h24 and h45 (**Figure 4H right**). Taken together, we propose that EME treatment blocks translocation by trapping the mRNA and eEF2 in place, which results in the formation of a more flexible, minimal contact disome.

### Collision structural features explain inhibitor-specific differences in activation of the RSR and ISR

Next, we aimed at understanding how the unique structural arrangement of EME-induced collisions may affect translational stress responses. Unlike EME treatment, ANS-induced collisions are a potent activator of the RSR due to their recognition by the kinase ZAK. The most notable structural difference between EME and ANS collisions is the translation state of the stalled ribosome. While the ZAK-bound ANS stalled (ZAK-ANS) 80S is observed in a classical POST state^19^, the EME stalled 80S occurs as a TI-4/TI-5 translocation intermediate. Although the 40S body is nearly identical between these states (∼1.6° rotation), the 40S head has yet to undergo the major ∼19° back-rotation to complete the translocation step (**Figure 5A**). This distinct rotation of the 40S head appears to profoundly influence the resulting collision interface. Compared to the EME disome, the collided 80S of the ZAK-ANS disome is tilted (∼9° rotation) and positioned closer to the stalled 80S (∼13 Å translational movement), resulting in a tighter overall collision (**Figure 5B**).

**Figure 5:**
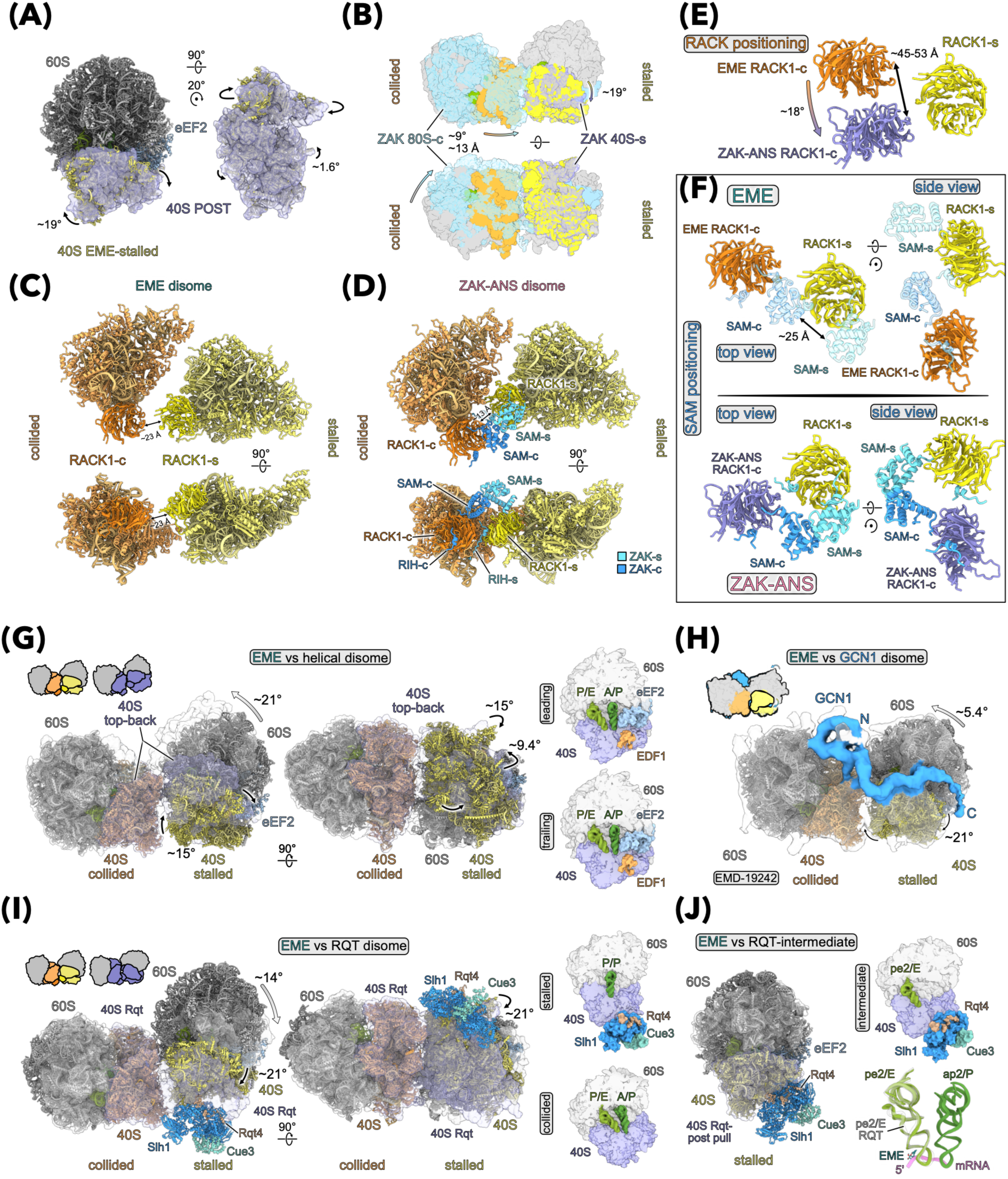
Structural differences between EME induced and other disome architectures. **(A)** Comparison of the EME stalled 80S and the classical POST state 80S focusing on the 40S conformation. The POST state 40S (purple) is shown as surfaces representation (PDB:9P72). Models are aligned by their large subunits. **(B)** Comparison of the EME and ZAK-ANS (PDB:9RPV) disome. Surface representations are shown and models were aligned by the large subunits of the stalled ribosomes. For the ZAK-ANS disome the collided (-c) 80S is shown in light blue and the stalled (-s) 40S in purple. Arrows indicate the directionality of movements with colors indicating start and end points. **(C and D)** 40S head-head interface for the EME (C) and the ZAK-ANS (D) disome. Distances measure the RACK1-s-RACK1-c separation in both disomes. **(E)** Comparison of the RACK1-s-RACK1-c positioning for the EME and ZAK-ANS disome. Distance measured is the distance range for all RACK1-c residues. Colored arrow indicates the rotational movement for RACK1-c relative to RACK1-s. Models are aligned to RACK1-s. **(F)** Comparison of the potential ZAK SAM domain positioning for the EME (top) and ZAK-ANS (bottom) disome. SAM domains for the EME RACK1 proteins (transparent) were imposed based on the ZAK-ANS disome (PDB:9RPV). **(G-J)** Comparison of the EME disome with helical disomes (G) (PDB:9B0Q), the GCN1 disome (H) (EMD-19242), and the yeast RQT disome (I) (PDB:7ZUW,7ZUX) and its intermediate state (J) (PDB:7ZRS). The EME disome or stalled 80S are shown as molecular models, and other disomes or 80S are shown as transparent surface representations. For the GCN1 disome, cryo-EM density is shown. All the disomes are aligned to the collided 80S. Cartoon representations are colored in the same way as the molecular models underneath. Cut-through views show the tRNA states and translation factors of the respective compared disomes. Black arrows indicate the respective 40S head or body rotations of the compared stalled 80S to the EME state. Colored arrows indicate the overall rotational difference of the stalled 80S with respect to the collided ribosome. The molecular overlay of tRNAs between the EME stalled 80S and the yeast RQT intermediate state are shown in (J).

A central requirement for ZAK activation is the dimerization of the ZAK SAM domains, which is driven by the RACK1-RACK1 (RACK1-s, RACK1-c, respectively) collision interface that critically positions the SAM domains of each ZAK monomer^19^. While the two 40S heads are notably separated in EME disomes, the distance between the RACK1 proteins is increased by only ∼10 Å (EME: ∼23 Å; ZAK-ANS: ∼13 Å), a gap that could possibly be overcome by the range of motion of the ZAK SAM domains (**Figures 5C,D**). However, comparing the ANS and EME disomes, the relative orientation of the RACK1s to one another differs strongly. Relative to RACK1-s, the RACK1-c is displaced by a rotation of ∼18° (∼45-53 Å distance) in the EME disome (**Figure 5E**). Due to this difference in RACK1-c positioning in the EME disome, the SAM domains of ZAK are situated too far from one another (∼25 Å) and are unsuitably oriented to form the head-to-tail dimer required for ZAK activation (**Figure 5F**). Therefore, the substantial difference we observe for RSR activation when comparing EME and ANS treatments is likely a consequence of the differences in the resulting disome architectures, and more specifically the distinct RACK1-s to RACK1-c orientations.

Similar to the lack of RSR activation, we also fail to observe an EME-induced increase in eIF2α phosphorylation and ISR activation (**Figure S6A**) in line with a recent report^32^. Recently, higher order helical polysomes that accumulate under stress have been structurally characterized in detail *in situ*^26,47^. Similar to EME collisions, their arrangement has been argued to be incompatible with ISR activation via the GCN pathway^26^. If we reduce the helical polysomes to a series of iterated disomes, the helical polysomes are comprised of stacked pre-translocation hybrid state ribosomes. As such, the collided 80S in the EME-stalled disome and the trailing ribosomes in helical polysomes are identical (**Figure 5G**). However, relative to helical polysomes, the EME-stalled 80S differs significantly for the small subunit (head: ∼15°; body: ∼9°), which results in a ∼21° rotation of the first ribosome with respect to the second (**Figure 5G**). Although helical collisions are structurally distinct from EME disomes, they both differ from GCN1-bound disomes observed in *in situ* cryo-ET data where the stalled ribosome is in the POST state and the collided ribosome is in the pre-translocation hybrid state^26^. When comparing these GCN1-bound disomes to our EME disomes, the main difference is the head rotation of the small subunit of the stalled 80S (∼21°), resulting in a different disome arrangement (∼5° rotation of the collided with respect to the collided 80S); these GCN1-bound disomes are most similar to the ZAK-ANS disome (**Figure 5H**). Importantly, GCN2 kinase activation has been argued to require an empty A site on the stalled 80S^17^, which here is blocked by the EME-trapped eEF2, and as recently noted^32^. While difficult to detect helical polysomes per se in sucrose gradients (**Figure 2A**), it is likely that under our standard EME treatment, helical polysomes form upstream of EME-stalled ribosomes due to inefficient stalling clearance by ASCC3 (**Figure 2**). Together, we suggest that EME-induced disomes and potentially helical polysomes are poor triggers for ISR and RSR activation because of their distinct structures relative to ANS-induced disomes.

### Collision structural features explain differences in clearance by ASCC3

Based on our biochemical data, the accumulation of ribosomes on mRNAs under EME treatment, but not ANS or DDB treatment, is a result of EME collisions resisting clearance by ASCC3 and the RQT (Ribosome Quality control-Trigger) pathway^48^. Together with observed increased eS10 ubiquitylation levels (**Figure 1A**), these data point towards the proper recognition of EME-induced collisions by ZNF598 but subsequent failure of downstream RQT activity. To understand the underlying cause of the persistence of EME stalls, we compared our EME disome with available structures of yeast RQT-bound disomes^9^. The RQT-bound disome consists of a canonical stalling combination of POST stalled and hybrid collided ribosomes (∼21° rotation of stalled 40S head), as seen with ANS-induced disomes; as discussed above, the collision interface is tighter with the stalled 80S rotated ∼14° towards the collided 80S (**Figure 5I**). Of note, RQT helicase activity pulls the mRNA in the opposite direction relative to translation, initially leading to a counterclockwise rotation of the head and simultaneous backtracking of the tRNAs from P/P, E/E to ap2/P and pe2/E. This RQT intermediate state effectively mirrors the EME-induced translocation intermediate (**Figure 5J**). As such, we infer from our structural data that there are several likely roadblocks for RQT-mediated splitting. First, the mRNA appears to be tightly locked in place by EME due to a network of interactions (**Figure 4H**). Second, the 5’-3’ directionality of the eEF2-driven mRNA translocation directly opposes the movement triggered by the 3’-5’ force applied by RQT helicase activity. Third, the conformation of the EME-stalled 80S already adopts a stabilized conformation similar to that observed after initial RQT activity. Finally, with the absence of a tight 40S head-head interface, persistent mRNA pulling would lack the wedge normally provided by the collided ribosome that likely contributes to the splitting of the stalled 80S^9^. Together, these structural observations nicely rationalize the complete lack of clearing of EME-induced disomes by RQT.

## Discussion

The ribotoxic stress response (RSR) was first defined by Iordanov *et al*.^25^ who noted that different translation inhibitors vary in their ability to activate the RSR. The mechanism behind these differences remained unclear even after the identification of ZAK as the key upstream MAP3K activator of the RSR^22,49^. Here we demonstrate that inhibitor-specific differences in RSR activation are due to differences in collision conformation. We further show that collisions induced by ANS, EME, and DDB differ in their sensitivity to collision-mediated quality control pathways. Overall, our work highlights the orientation of the collision interface as a key determinant of RSR activation while clarifying important properties of drugs commonly used to study collision-mediating signaling. Importantly, despite all being commonly used elongation inhibitors^50^, ANS, EME, and DDB exhibit distinct effects on translation and quality control that warrant careful consideration when used to study collision-mediated signaling.

We utilized RNA-seq to dissect the contributions of inhibitor-specific ZAK-dependent and ZAK-independent signaling on early transcriptome changes. In particular, we show that the subset of transcripts induced by the ZAK-mediated RSR are those most highly expressed under collision-inducing conditions. These observations are consistent with our previous study that defined the RSR as the key driver of early phosphoproteomic changes downstream of UV irradiation^16^. Many of the transcripts we identify as ZAK-dependent were also identified in a previous study that showed the *Legionella pneumophila* effector SidI can stall ribosomes and induce RSR activation, highlighting the importance of ribosome collisions in contexts beyond elongation inhibitor treatment^51^. Other bacterial toxins targeting the ribosome, such as diphtheria toxin, also potently activate the RSR^25,52^. Additionally, our finding that elongation inhibition activates ZAK-independent transcriptional programs, such as that induced by *MLXIP*, is in agreement with previous studies showing that general translational inhibition activates transcriptional programs^37,53^; it is not surprising that the energetically consuming process of translation determines critical cellular responses. We also highlight that other pathways, such as the one driving *HES1* expression, contribute to our observed transcriptional responses but remain uncharacterized.

Aside from each drug’s ability to induce the RSR (and the ISR), we considered other inhibitor-specific effects on collision clearance. In addition to being highly tolerant to flexibility of the collision interface^7^, we find that ZNF598 is remarkably quick in identifying collisions (**Figure S7B**), in line with its known role in negatively regulating collision-mediated signaling^17,18,23^. We further find that while ANS- and DDB-induced collisions are cleared by ASCC3, EME-induced collisions are not (**Figure 2B**). We note that contrary to our observations, ASCC3-mediated clearance of EME collisions has been previously reported^23^; this study utilized a transient EME treatment prior to drug washout which may explain the differences in our results. Our data indicate that a key marker of ASCC3 activity is an accumulation of 80S complexes on polysome profiles as we observed with ANS and DDB treatment, but not EME (**Figure 2**). A recent cryo-ET study of ANS-treated cells suggested that the 80S accumulation that they observed reflected the ZNF598-mediated clearing of the lead ribosome in a collision queue^26^; our polysome data are nicely consistent with their proposed model where the 60S and 80S peaks seen with ANS treatment are dependent on the presence of ASCC3.

We also noted ANS-specific accumulation of 60S subunits and a sharp decrease in polysomes over time. These observations are supported by early studies of ANS, which ascribe ANS-mediated 60S accumulation to a defect in 60S joining during initiation^54^. This mechanism would also rationalize the ANS-induced decrease in polysomes over time (**Figure 2A**) and the loss of RSR activation at long timepoints (**Figure 1A**); once collided ribosomes are cleared by ASCC3, initiation is inhibited and thus no more collisions can accumulate. Importantly, the large accumulation of 80S ribosomes downstream of ANS treatment appears to be independent of the ISR. The ANS-induced eIF2α phosphorylation previously reported by our lab^14^ is likely modest and does not yield a major phenotype in polysome profiles reflecting the ISR^39^ as compared to true ISR agonists like UV and amino acid starvation^16,32^ (**Figure 2C**). We note that our new data here agree with a previous study in yeast proposing that GCN2 activation is triggered by stalled (or colliding) ribosomes with an unoccupied A site^17^; in this case, ANS and DDB likely lead to the formation of ribosome states with mostly, but not completely, occupied A sites^28,55^. In contrast, more potent activators of GCN2 such as UV, MMS or amino acid starvation leave the A site more fully unoccupied^16,17,32,56^.

Recent work from our labs elucidated the mechanism of ZAK activation at the ANS-induced disome interface, where the RACK1 proteins of the collided ribosome facilitate the dimerization of ZAK through its SAM domains and its subsequent autophosphorylation^19^. However, several independent studies have suggested that collisions may exist in distinct structural states^26,47^, which may favor or disfavor ZAK activation. In agreement with this, we observe that EME-induced disomes, which do not activate the RSR, are notably more loose and flexible compared to ANS-induced disomes, which do activate ZAK and are considerably more rigid^19^. Thus, we define disome geometry, particularly RACK1-RACK1 spacing, as the key determinant of ZAK activation (**Figure 6**). This parameter may allow the cell to discern pathological from non-pathological collisions. For instance, several studies have argued that collisions involving >3 ribosomes cannot exist in the canonical ZAK-activating disome interface (**Figure 6, left**); instead, these collisions adopt a helical polysome structure^26,47^. Helical polysome structures have been observed *in vitro* and *in vivo* across numerous studies^26,47,57–64^. The ribosome interfaces in this helical structure exhibit much greater RACK1-RACK1 distances that would disfavor ZAK activation and therefore prevent aberrant signaling being generated by heavily translated mRNAs (**Figure 6, right)**. We hypothesize that EME treatment induces formation of these helical assemblies given that EME-induced collisions are not cleared by ASCC3 (**Figure 2B**) and that the EME-induced increase in polysomes over time suggests that translation initiation is not blocked (**Figure 2A, middle**); indeed, there is essentially no ZAK activation in this context. We speculate that these limitations on RSR and ISR activation, in addition to the limitations inevitably imposed by kinetics, could be critical in cells where translation is robust and where collisions must regularly occur.

**Figure 6:**
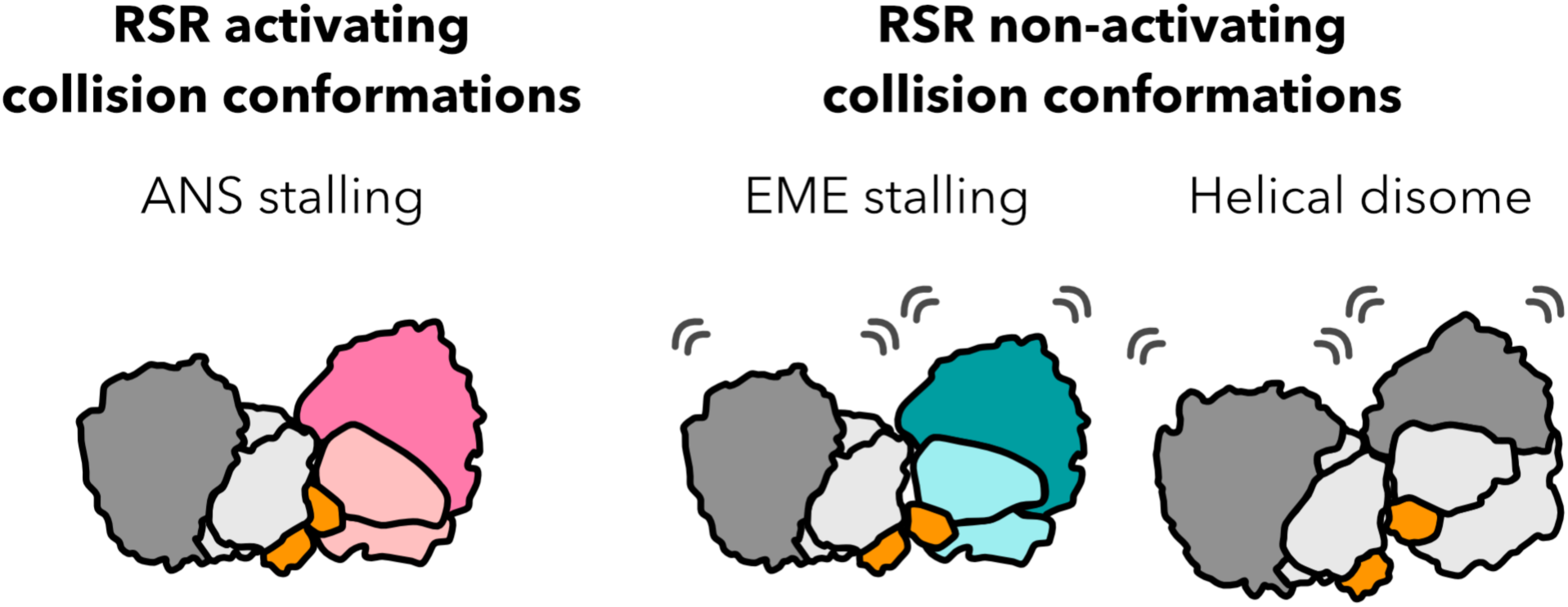
Model. **(Left)** Ribosome collisions formed after certain stalling events, such as ANS treatment, exhibit a rigid collision conformation with proximal RACK1 spacing that allows ZAK activation. **(Right)** Ribosome collisions formed after EME treatment or helical polysomes exhibit more flexible collision conformations with well-separated RACK1 spacing that disfavors ZAK activation.

Interestingly, these structural constraints that limit collision-mediated signaling activation may have broader implications in cells where translation is regulated at the level of elongation. In neurons, for example, ribosomes are paused during elongation in polysomes to facilitate rapid local translation in response to synaptic activities^65^. Ribosomes in these stalled polysomes are tightly packed and have been proposed to be stalled in a pre-translocation state though these cells show no evidence of collision-related QC pathway or signaling activation^66,67^. We speculate that such non-pathological neuronal stalled ribosomes may represent a situation in which the cell maintains stalled ribosomes without activating cellular stress responses.

## Supplemental Figure Legends

**Figure S1:**
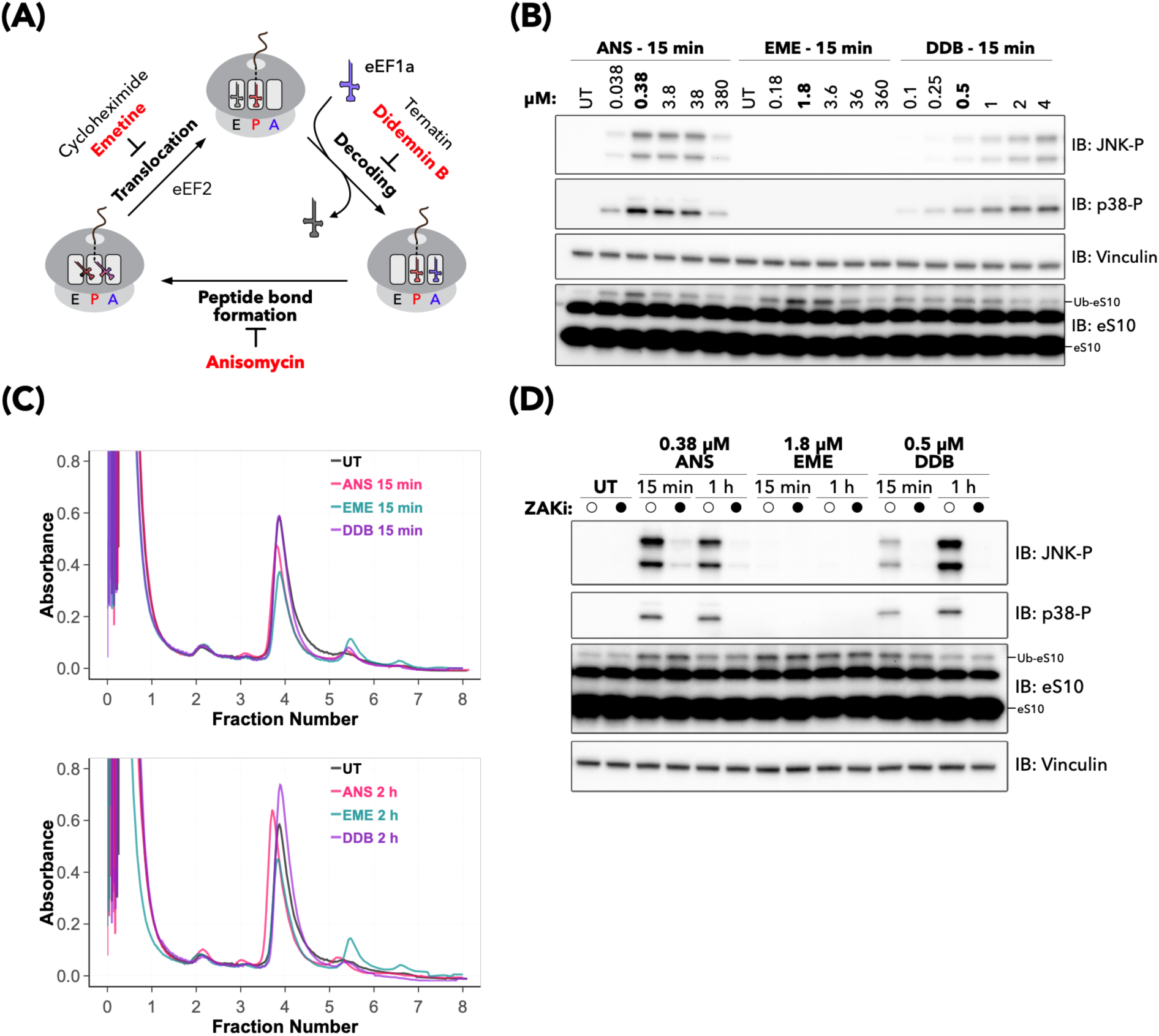
Characterization of elongation inhibitors used to induce the collision-mediated ribotoxic stress response (related to Figure 1). **(A)** Simplified overview of translation elongation and inhibitors targeting each step. Inhibitors used in this study are highlighted in red. **(B)** Immunoblots of HEK293T cells treated with varying doses of anisomycin (ANS), emetine (EME), or didemnin B (DDB) for 15 min. Doses used in the rest of the study are in bold. **(C)** Polysome traces of RNase A-treated lysates of HEK293T cells treated with 0.38 µM ANS, 1.8 µM EME, or 0.5 µM DDB for either 15 min (top) or 2 h (bottom). **(D)** Immunoblots of HEK293T cells pre-treated with DMSO or ZAK inhibitor (ZAKi) for 1 h prior to treatment with the indicated elongation inhibitor for the indicated times.

**Figure S2:**
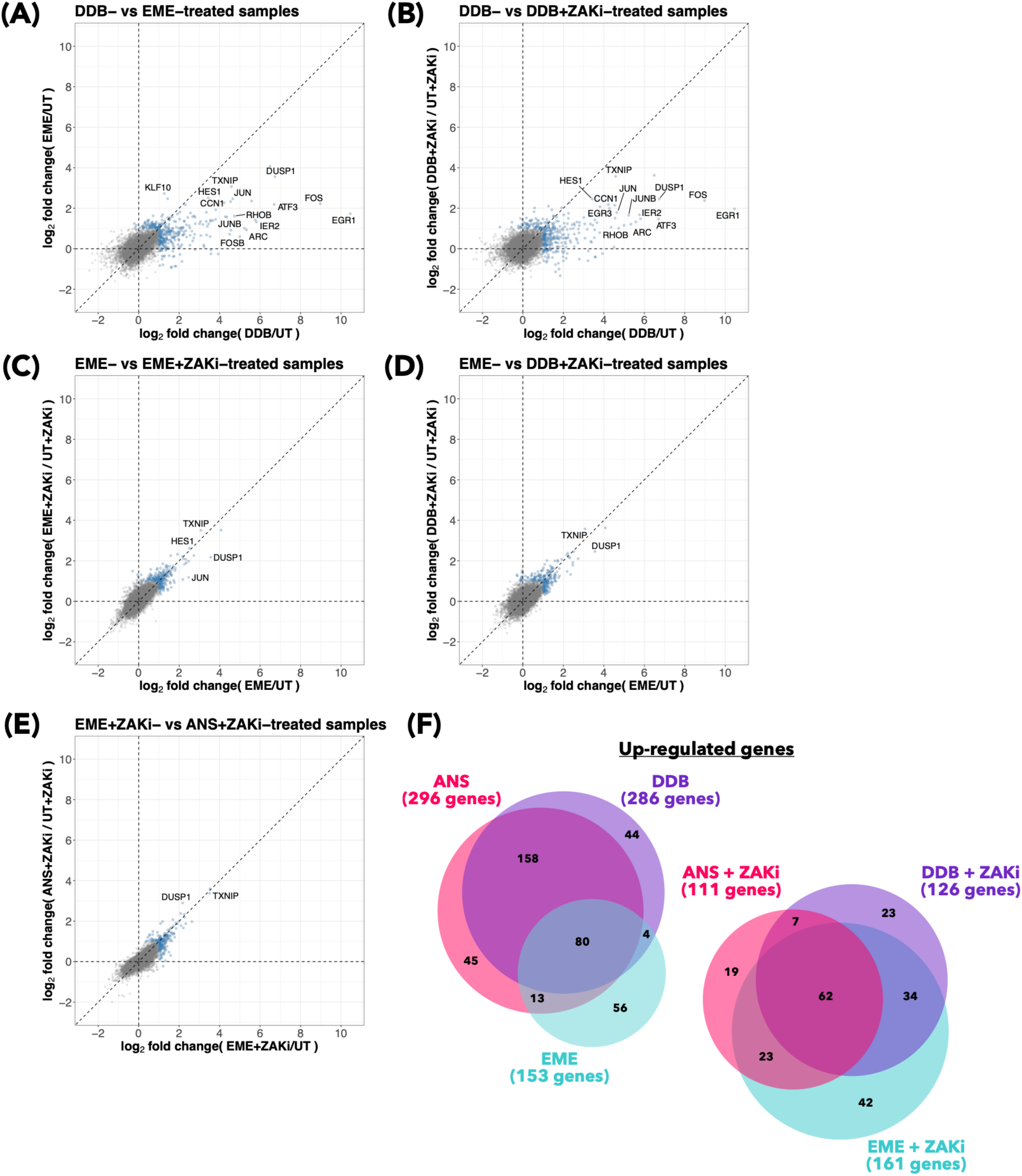
Comparison of gene expression changes induced by different elongation inhibitors measured by RNA-seq (related to Figure 1). **(A-E)** Cross-correlation plots of gene expression in the indicated inhibitor-treatment conditions normalized to the untreated condition. Points highlighted in blue indicate genes showing a log_2_(fold change) (log_2_FC) greater than 1 in at least one condition. **(F)** Area-proportional Venn diagram of significantly up-regulated genes (log_2_(fold change) > 1 & adjusted p-value (padj) < 0.01) in the indicated conditions.

**Figure S3:**
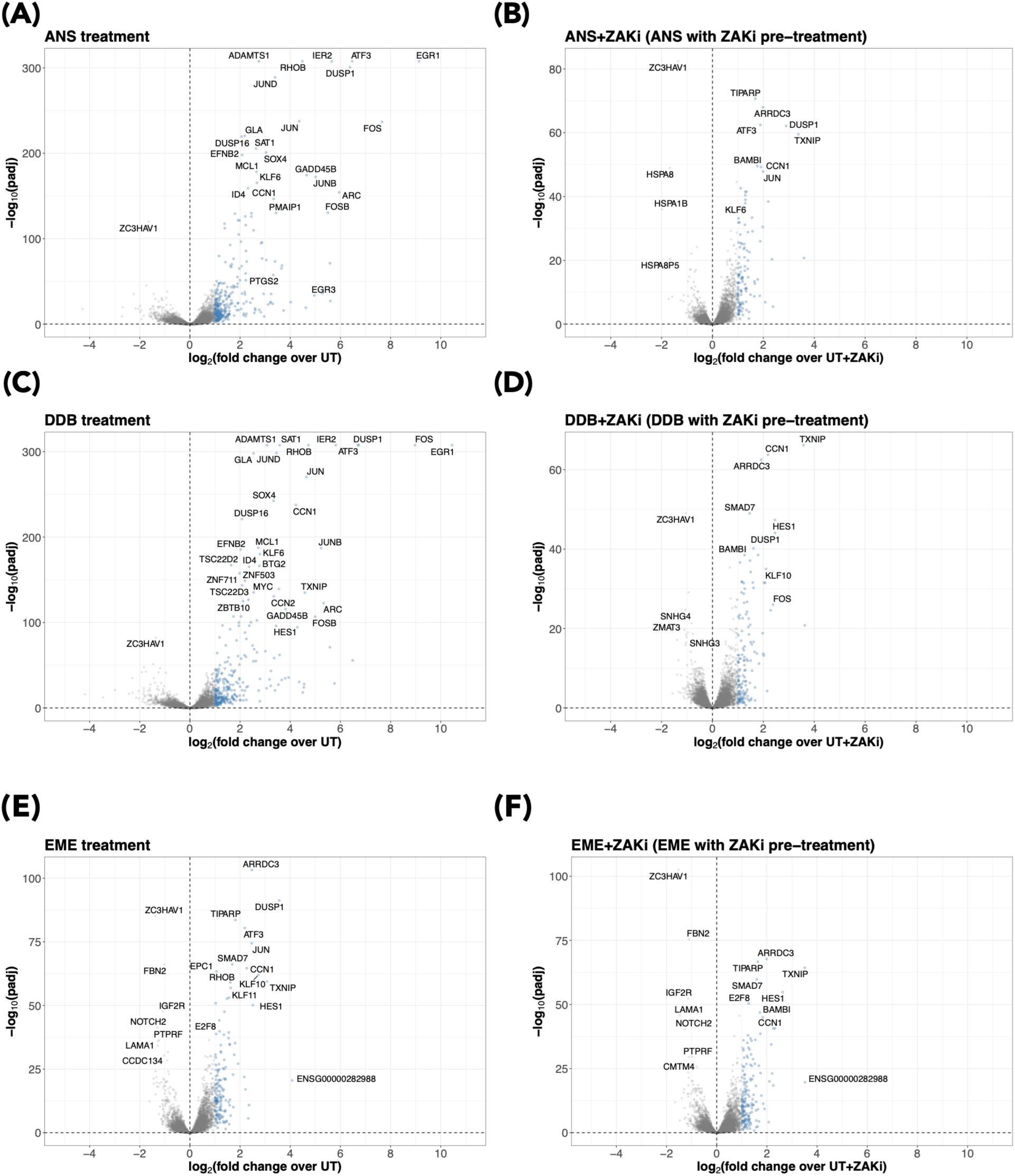
Gene expression changes induced by different elongation inhibitors measured by RNA-seq (related to Figure 1). **(A-F)** Volcano plots of differential gene expression in cells pre-treated with DMSO or ZAK inhibitor prior to treatment with the indicated elongation inhibitor, normalized to the untreated condition as measured by RNA-seq. Points highlighted in blue indicate genes showing a log_2_(fold change) greater than 1.

**Figure S4:**
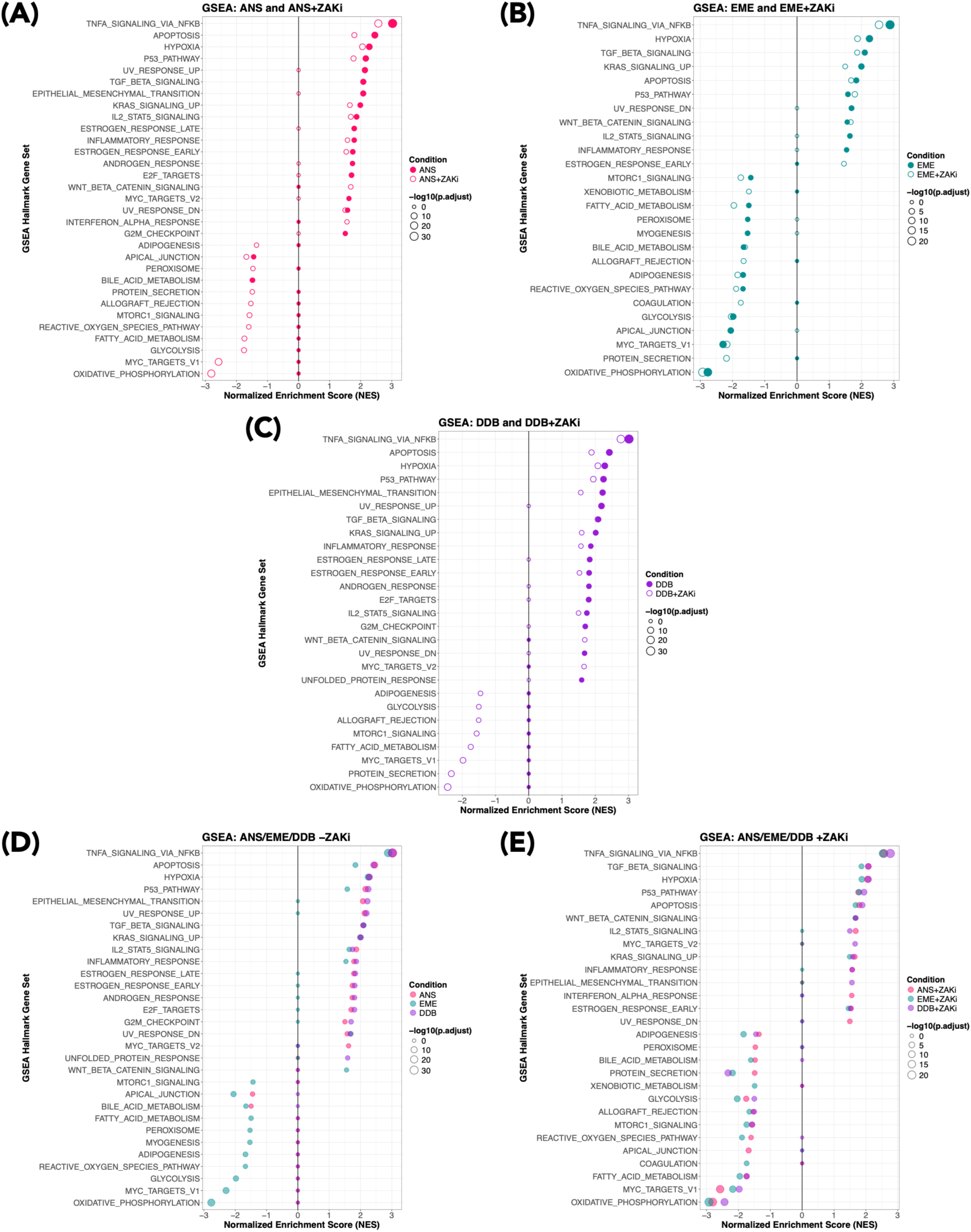
GSEA of gene expression changes induced by different elongation inhibitors measured by RNA-seq (related to Figure 1) **(A-C)** Dot plots of normalized enrichment scores (NES) for human MSigDB hallmark gene sets for the indicated drug and drug+ZAK inhibitor conditions. Solid dots correspond to drug alone, whereas hollow dots correspond to drug + ZAK inhibitor pre-treatment conditions. The size of the point corresponds to the -log10(adjusted p-value). (D) Dot plot of normalized enrichment scores (NES) for human MSigDB hallmark gene sets for all drug (without ZAK inhibitor pre-treatment) conditions overlaid. The size of the point corresponds to the -log10(adjusted p-value). **(E)** Same as D, but for the drug + ZAK inhibitor pre-treatment conditions.

**Figure S5:**
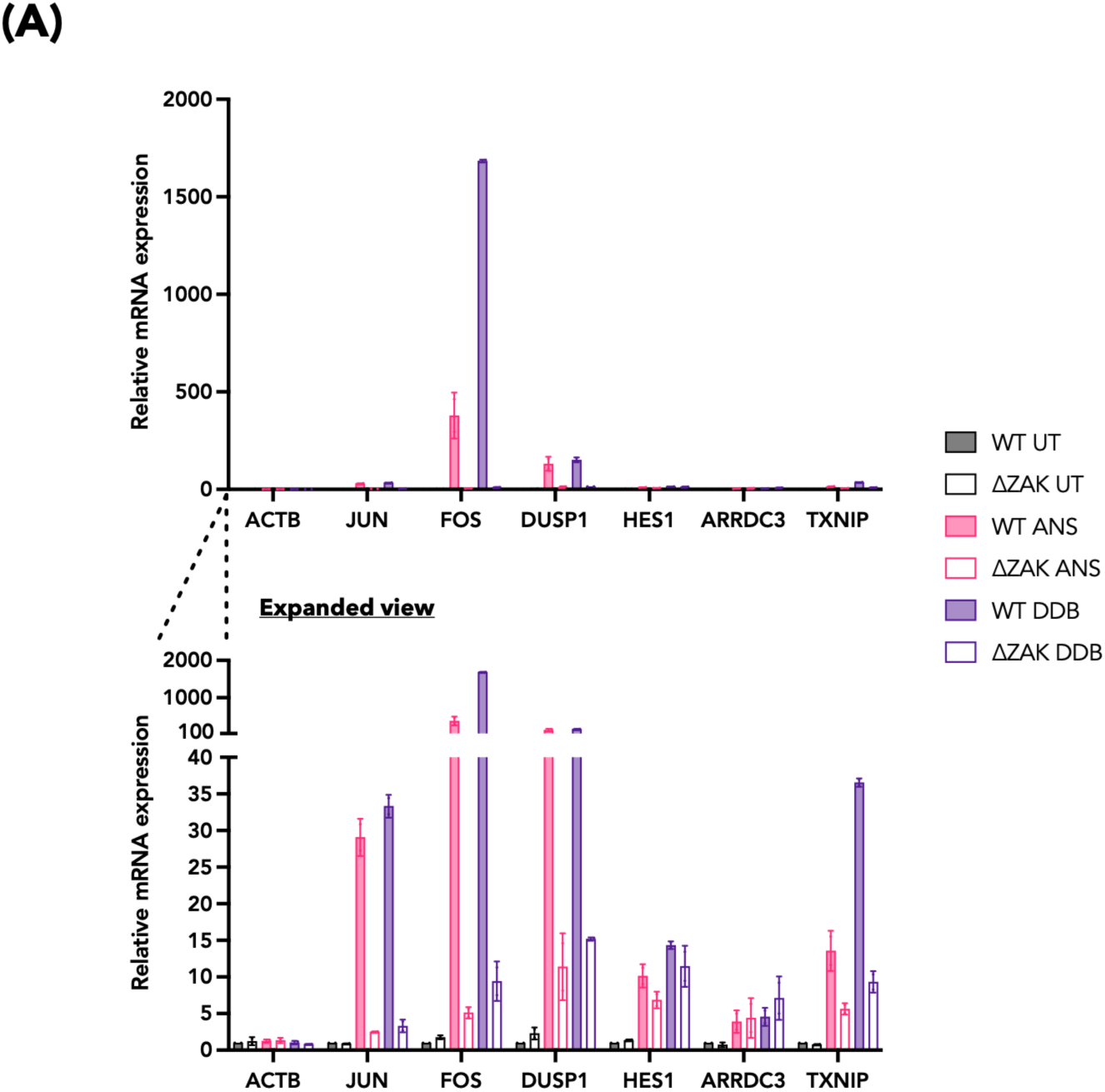
Validation of ZAK-dependent and ZAK-independent transcript induction (related to Figure 1). **(A)** RT-qPCR of wildtype (WT) or ZAK knockout (ΔZAK) HEK293T cells left untreated or treated with 0.38 µM anisomycin (ANS) or 0.5 µM didemnin B (DDB) for 2 h. The bottom plot shows the same data as the top plot, but with a smaller axis to visualize smaller changes in expression.

**Figure S6:**
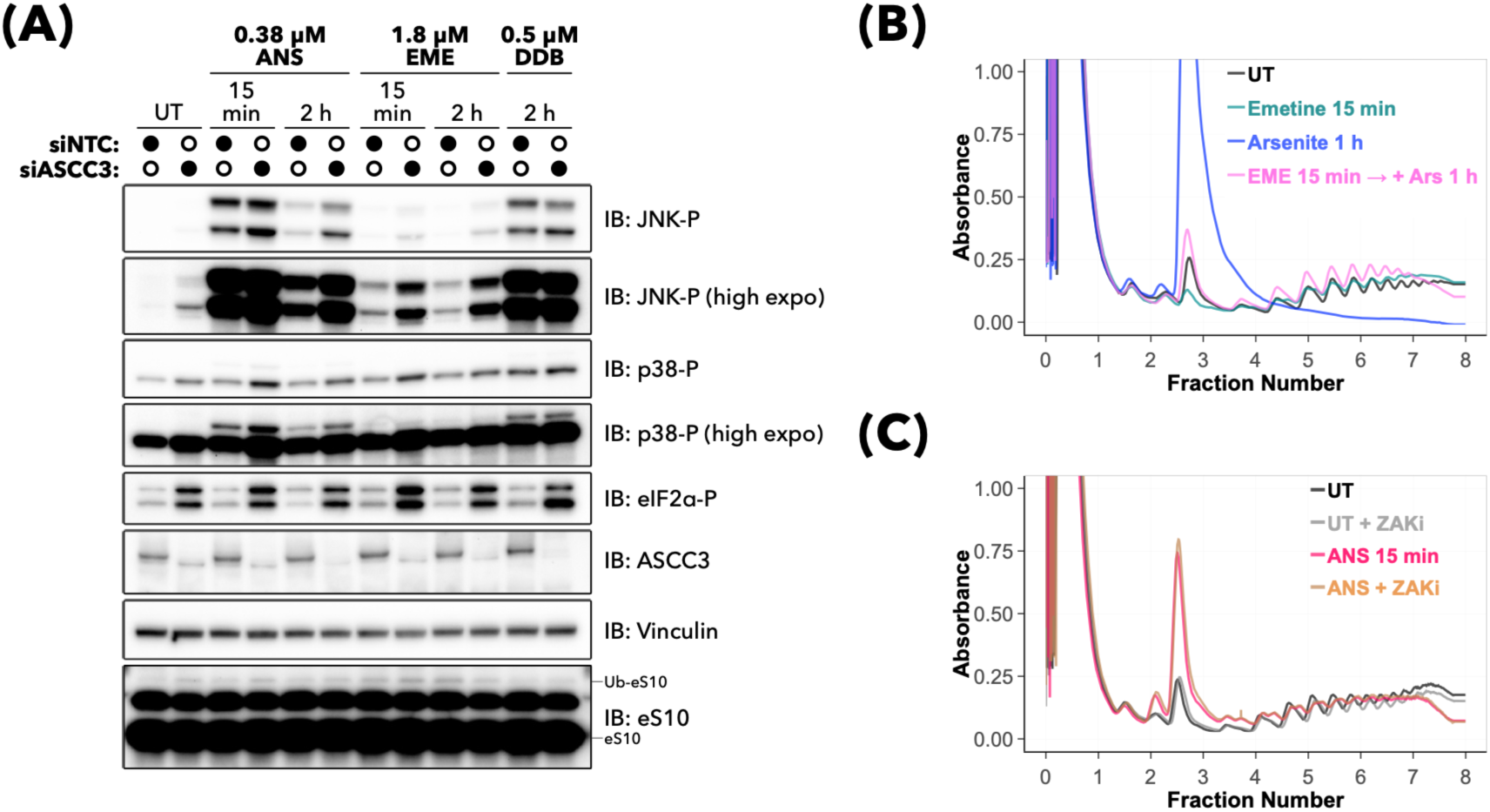
Characterization of ASCC3 activity and translation initiation on inhibitor-induced collision signaling and clearance. **(A)** Immunoblots of HEK293T cells transfected with non-targeting (siNTC) or ASCC3-targeting (siASCC3) siRNAs prior to treatment with the indicated doses and times of elongation inhibitor. **(B)** Polysome traces of HEK293T cells either left untreated (UT), treated with 1.8 µM emetine for 15 min, 100 µM sodium arsenite for 1 h, or 1.8 µM emetine (EME) for 15 min prior to treatment with 100 µM sodium arsenite (Ars) for 1 h. **(C)** Polysome traces of HEK293T cells left untreated or pre-treated with ZAK inhibitor (ZAKi) for 1 h prior to treatment with 0.38 µM anisomycin for 15 min.

**Figure S7:**
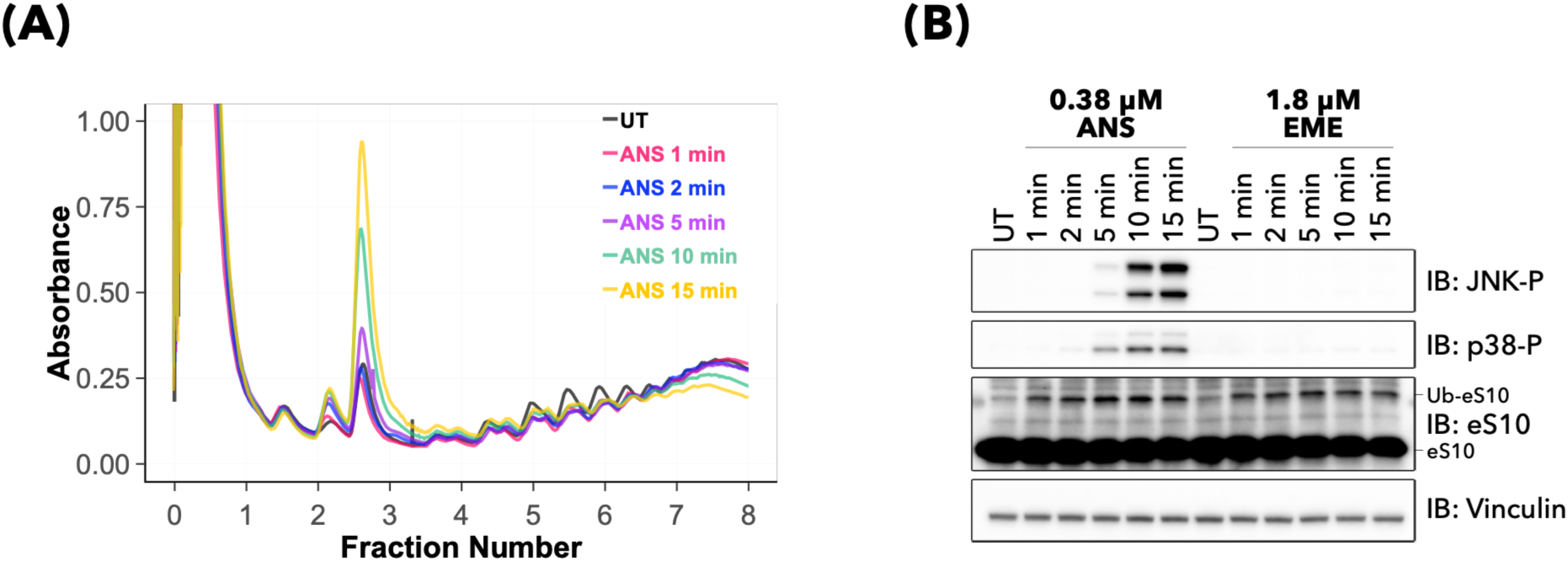
Characterization of elongation inhibitor effects at early timepoints. **(A)** Polysome traces of HEK293T cells treated with 0.38 µM anisomycin (ANS) for the indicated times. **(B)** Immunoblots of HEK293T cells left untreated (UT) or treated with the indicated dose and time of elongation inhibitor.

**Figure S8:**
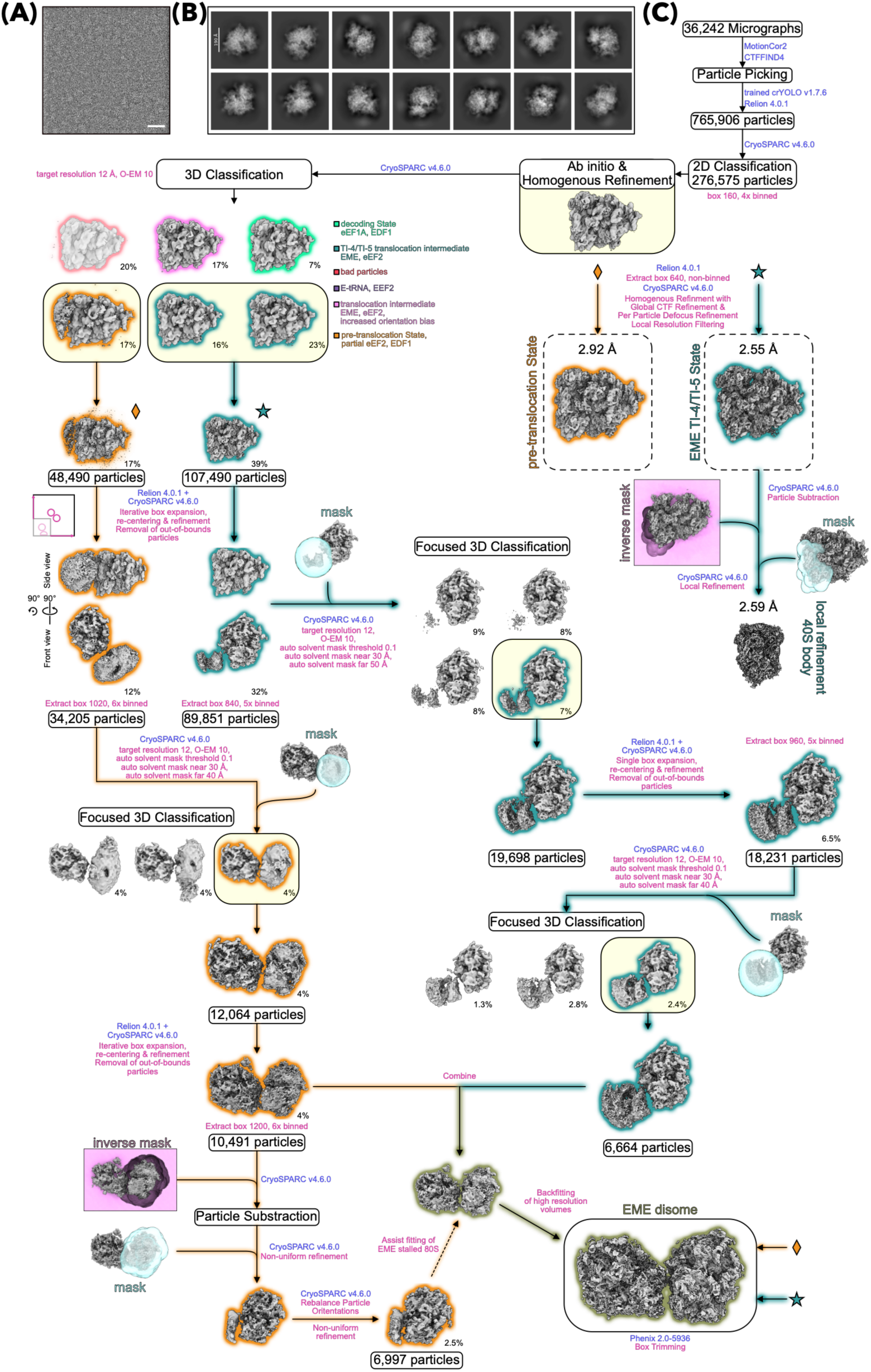
Data processing for the EME *in extracto* dataset. **(A)** Representative micrograph with inverted contrast and 20 Å low-pass filter applied. Scale bar corresponds to 35 nm. **(B)** Representative 2D class averages for ribosomes. **(C)** Cryo-EM data processing and sorting scheme. Selected classes from 3D classifications are framed. Relevant processing steps are highlighted, parameters are noted in pink, and software used in blue. Processing steps for the stalled 80S are highlighted in dark green and for collided 80S in orange. The orange diamond and the dark green star indicate intermediate particles for stalled and collided 80S, respectively, where processing is picked up elsewhere in the scheme. All relevant masks used are shown.

**Figure S9:**
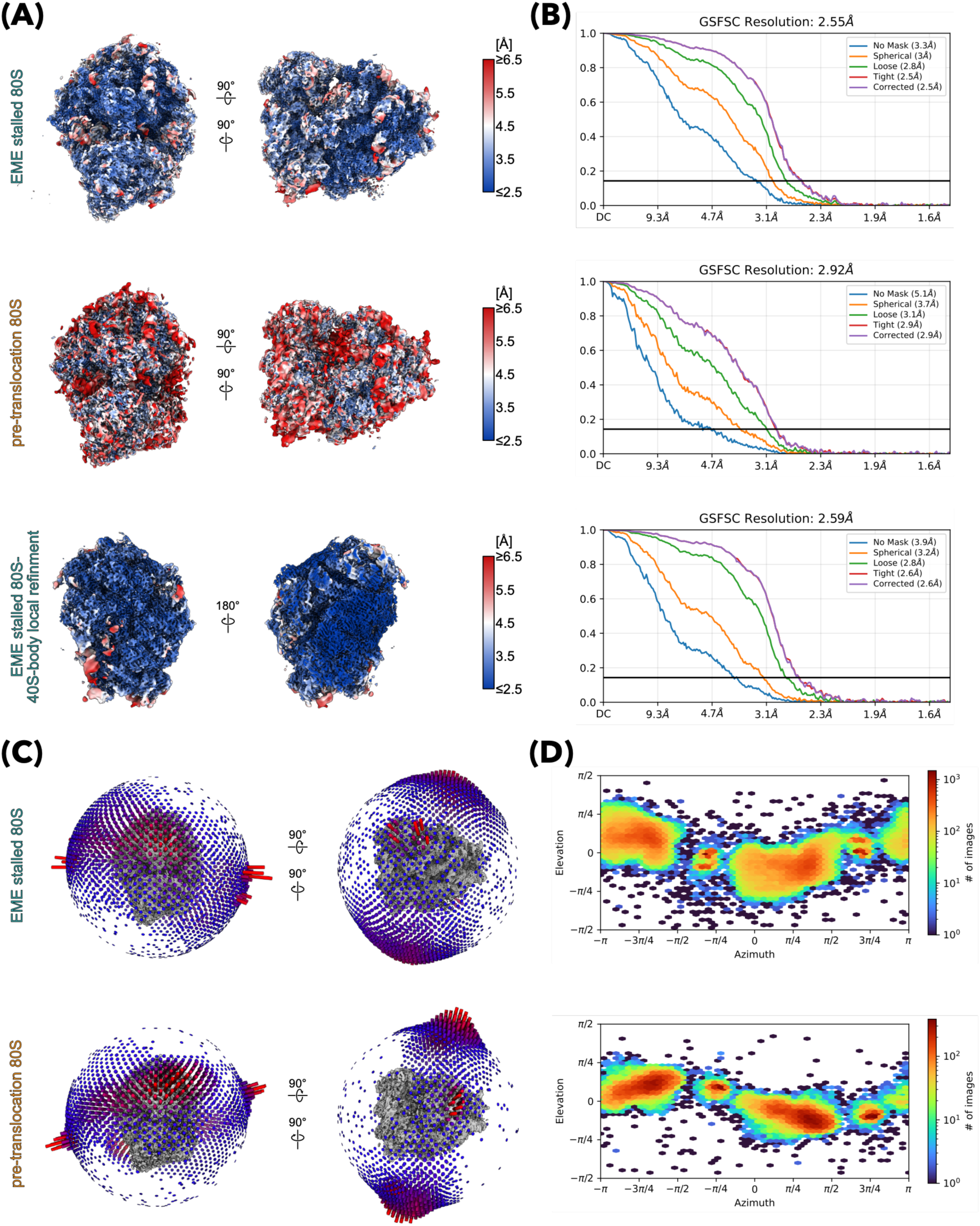
Cryo-EM reconstruction validation. **(A)** Cryo-EM maps filtered and colored according to local resolution. **(B)** Gold standard Fourier shell correlation curves (GSFSC, as generated by cryoSPARC) with 0.143 criterion for overall resolution estimation. **(C)** 3D representations of the orientation distributions of final particles. **(D)** 2D heatmap of particle viewing directions and numbers as generated by cryoSPARC.

**Table S1:**
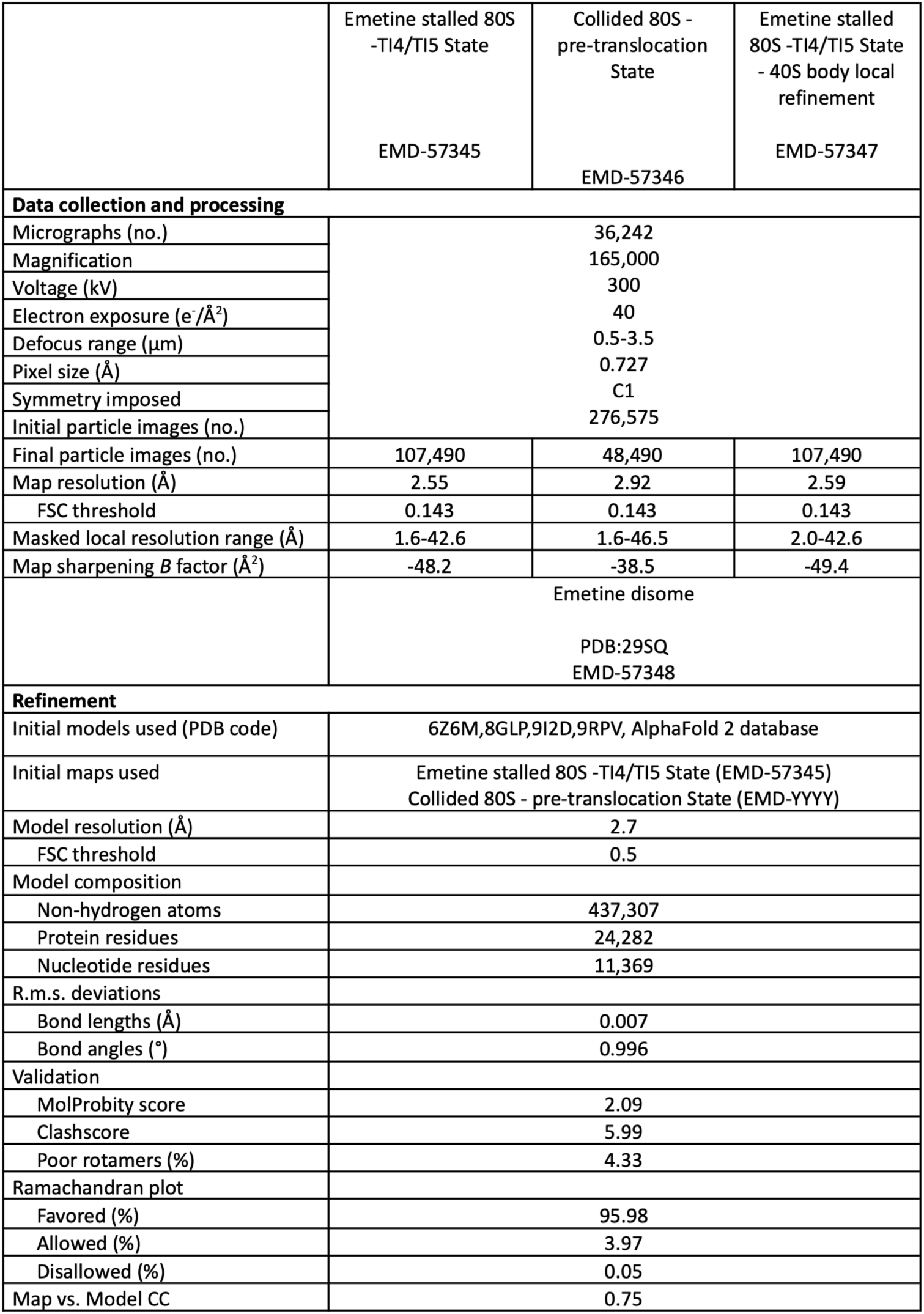
Data collection details and model refinement statistics.

## Acknowledgements

We thank Vienna Huso and Marco Catipovic for providing the ΔZAK HEK293T cell line, and Nicolle Rosa-Mercado for providing technical and computational assistance with RNA-sequencing and RT-qPCR. We thank Charlotte Ungewickel, Susanne Rieder and Otto Berninghausen for technical assistance with cryo-EM data acquisition. This study was supported by grants from the DFG (BE1814/20-1, BE1814/22-1) to R.B.; T.D. is supported by the Graduate School of Quantitative Biosciences Munich (QBM). A.R.B. is supported by NIH grant R01GM136960. R.G. is supported by the Howard Hughes Medical Institute (HHMI). Work in this paper was additionally supported by NIH grants R37GM059425 and R35GM156244 awarded to R.G..

## Author contributions

J.J.L., A.R.B., R.B., and R.G. conceptualized the study. J.J.L. conducted the biochemical and RT-qPCR assays, as well as RNA-sequencing library preparation and data analysis. Cryo-EM samples were prepared by T.D., cryo-EM data was processed by T.D. and interpreted by T.D. and R.B. with the help of M.T.. J.J.L., T.D., A.R.B., R.B., and R.G. wrote the manuscript. All authors reviewed and approved the manuscript.

## Materials and Methods

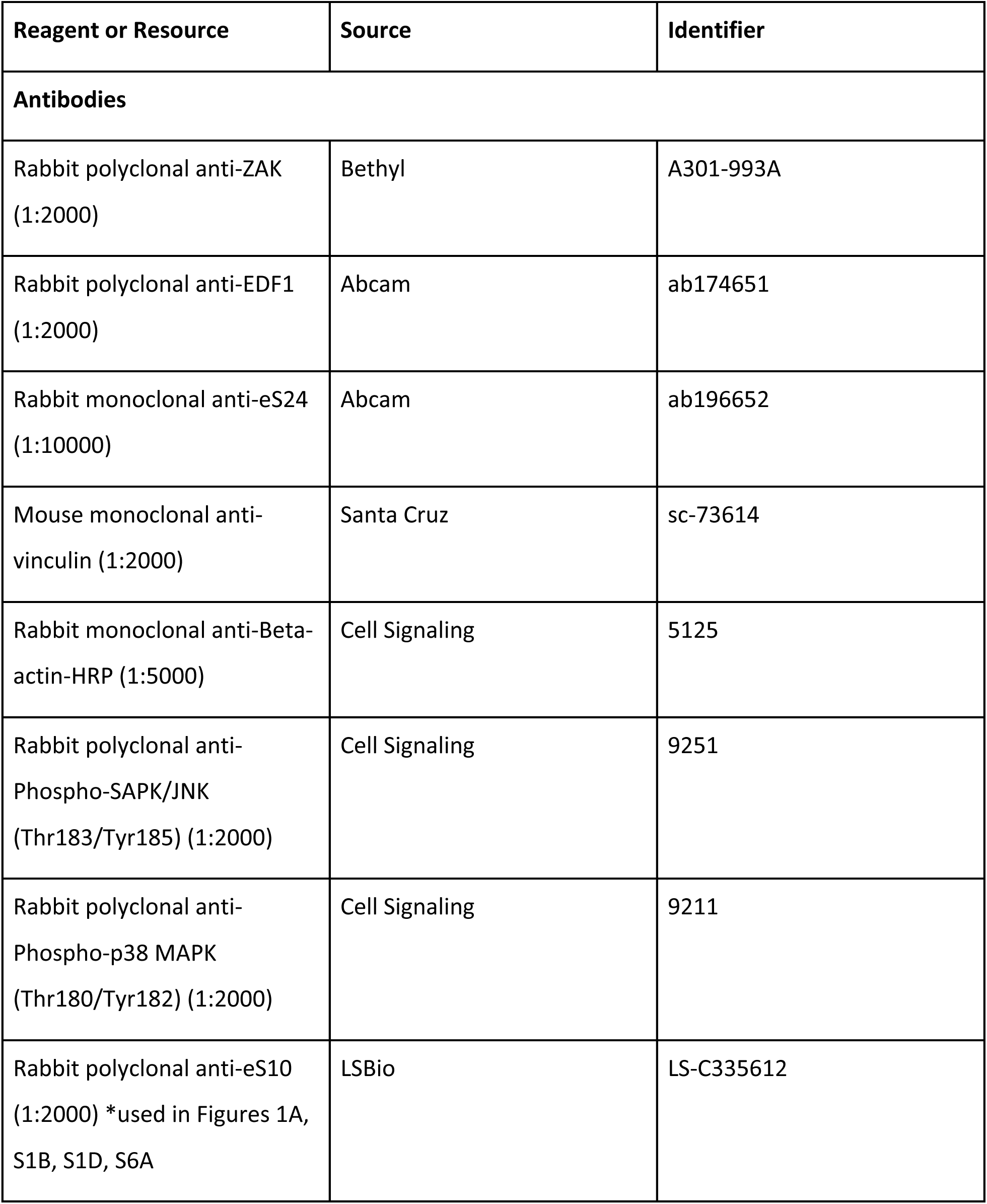
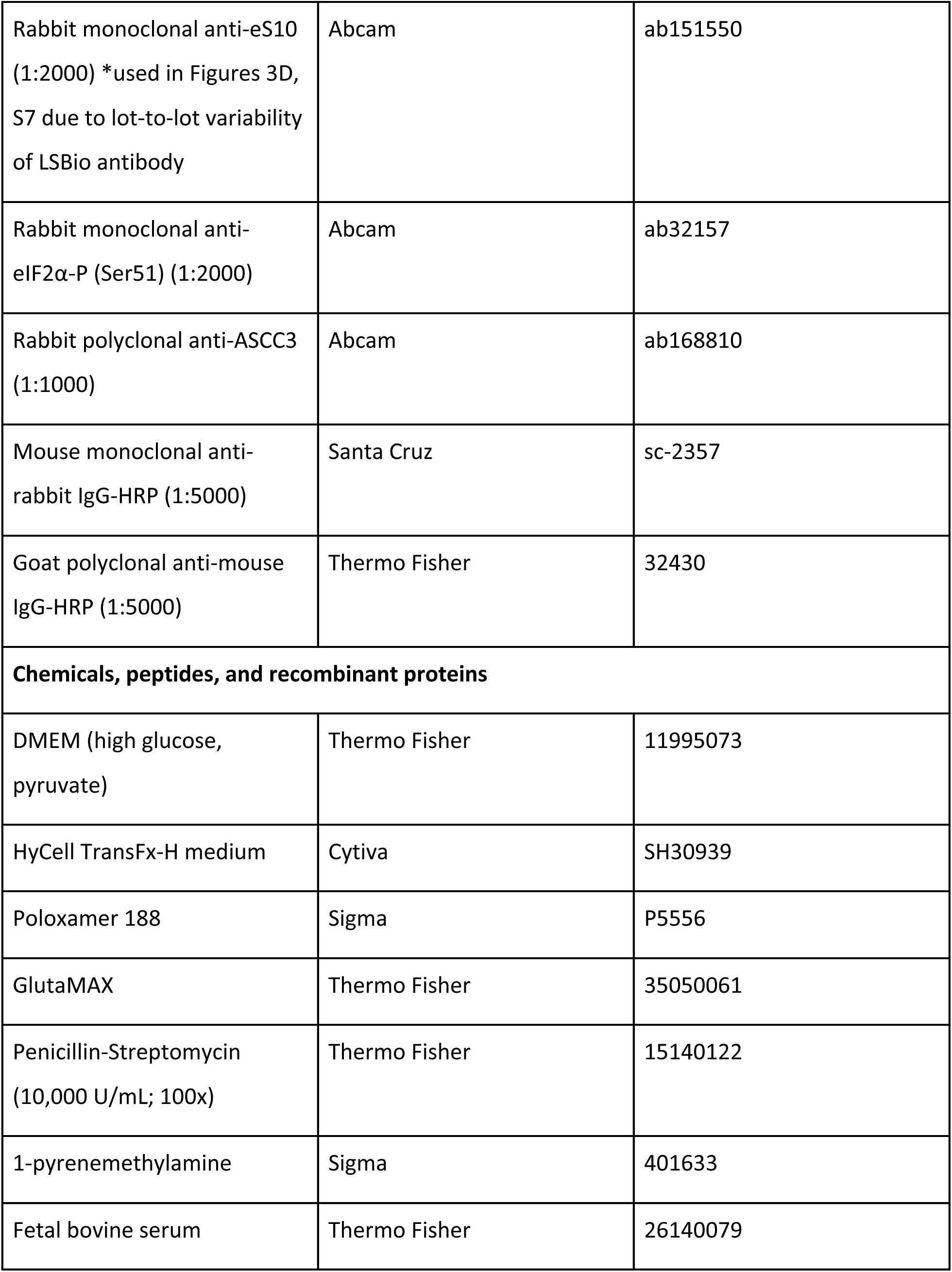
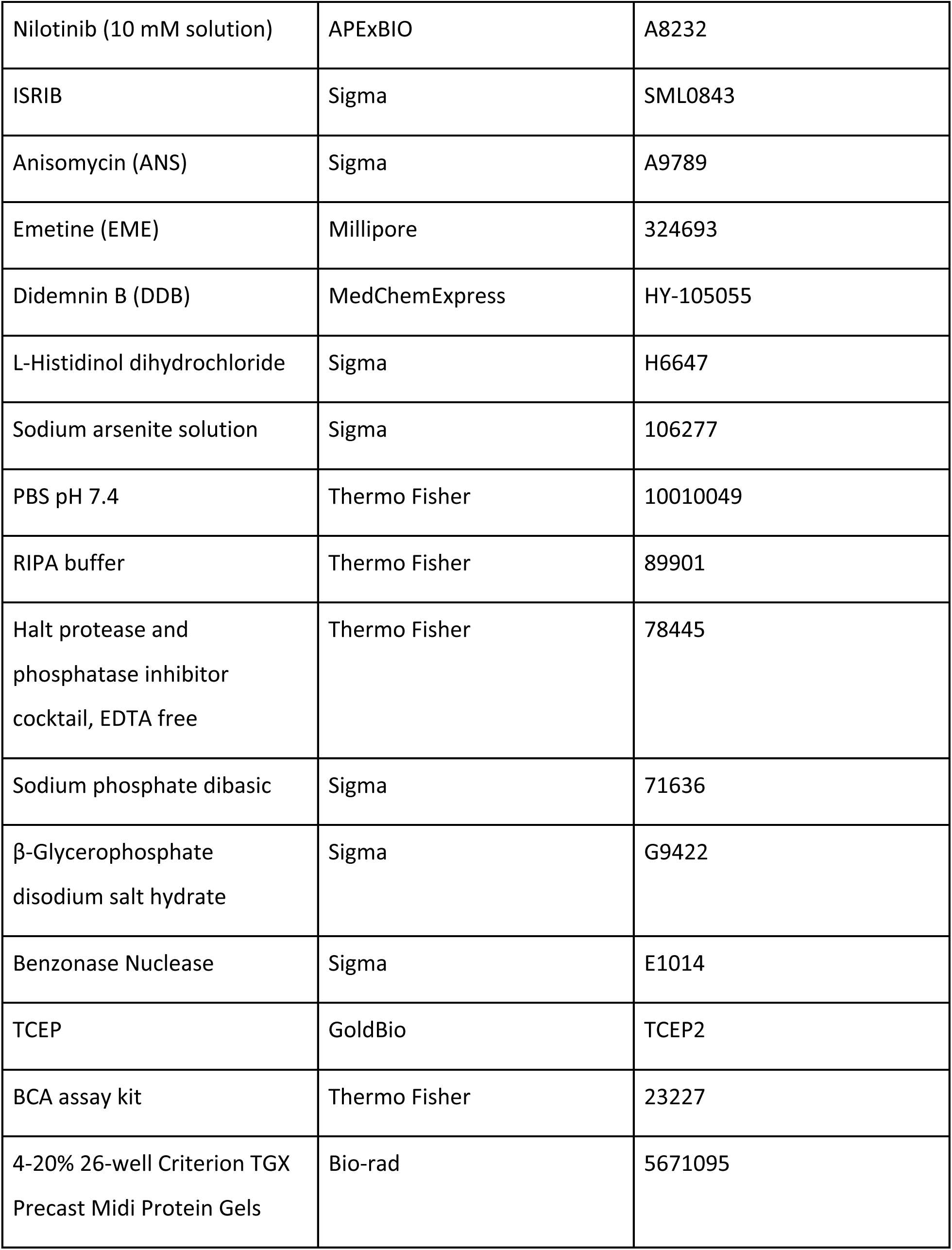
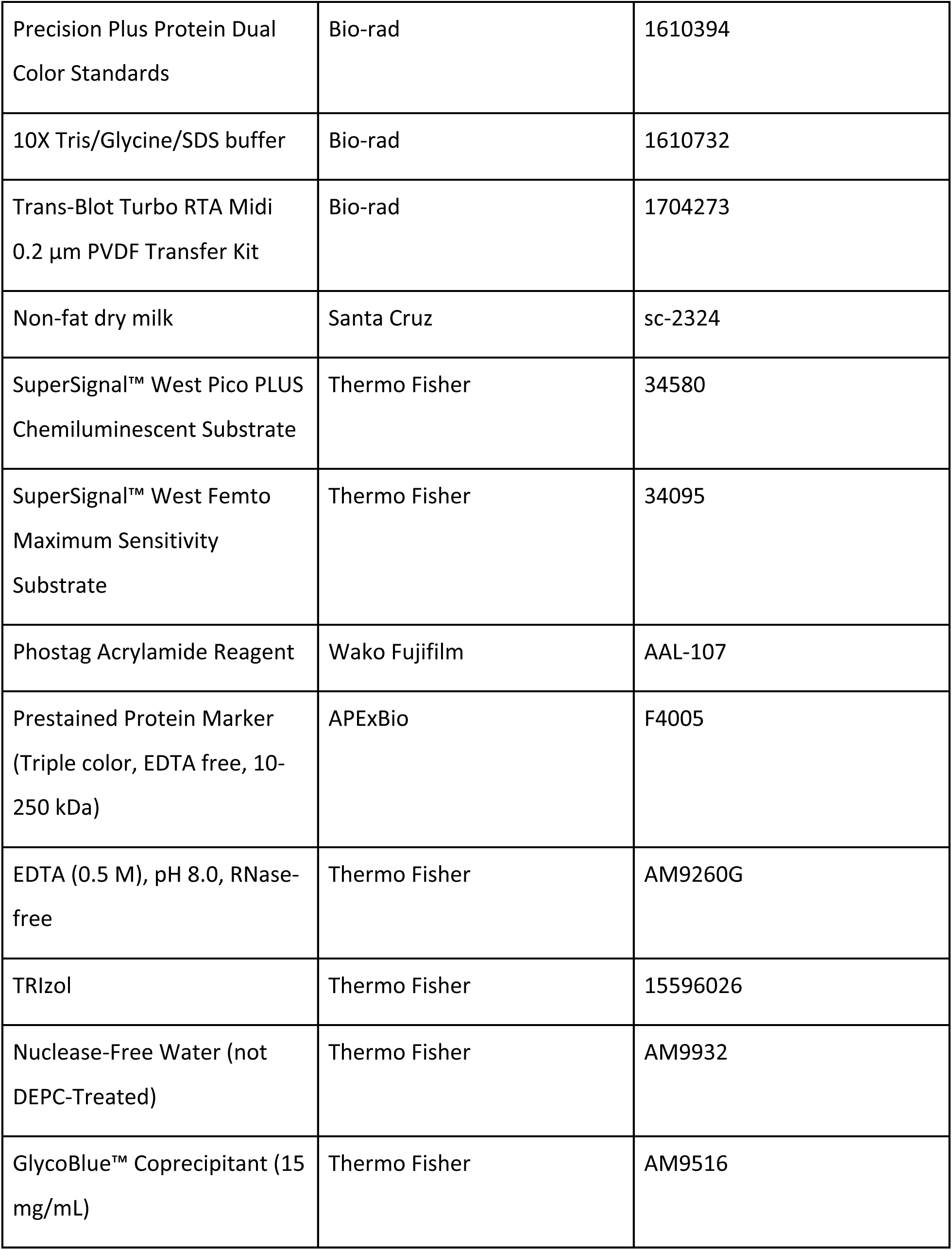
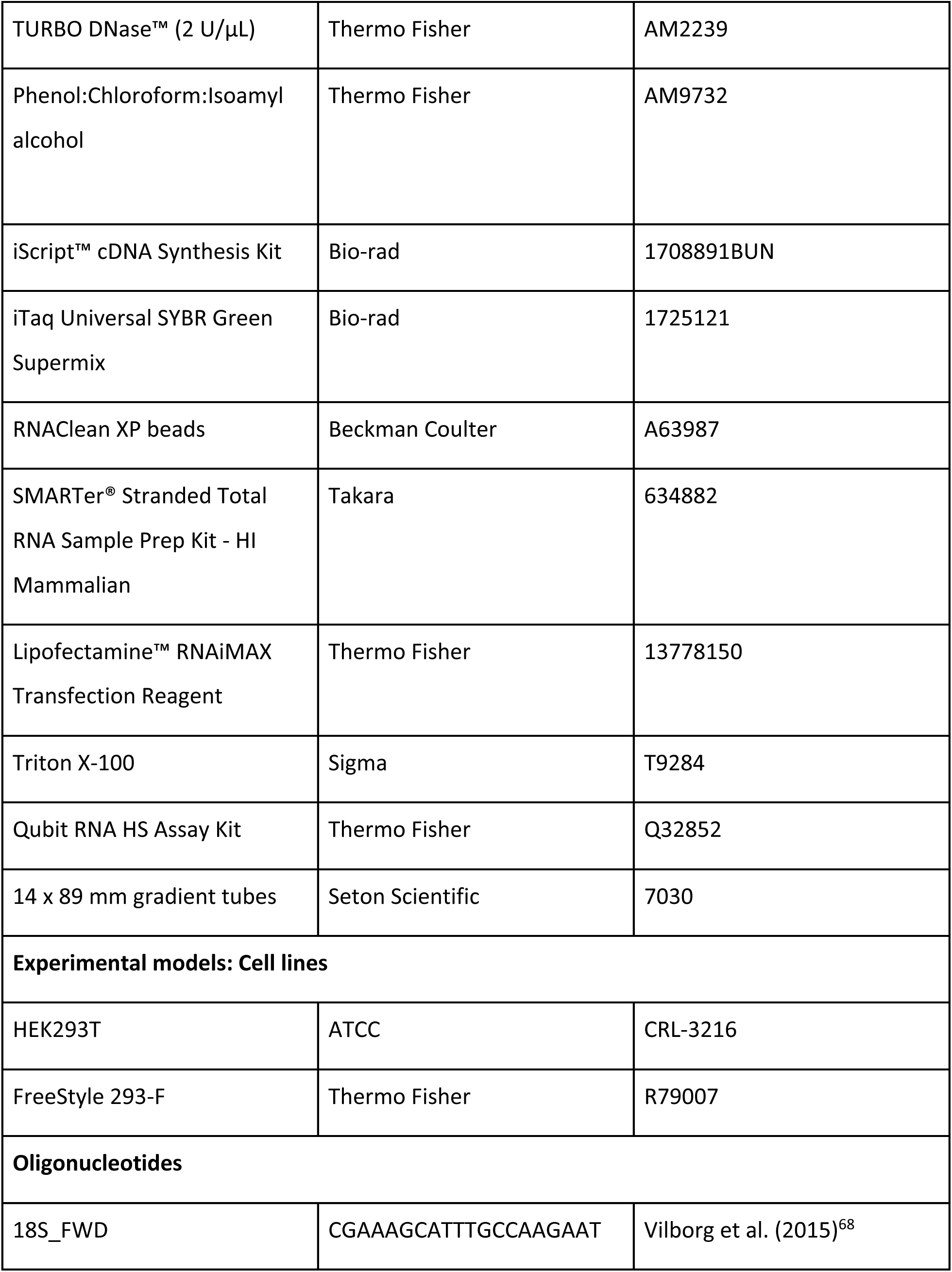
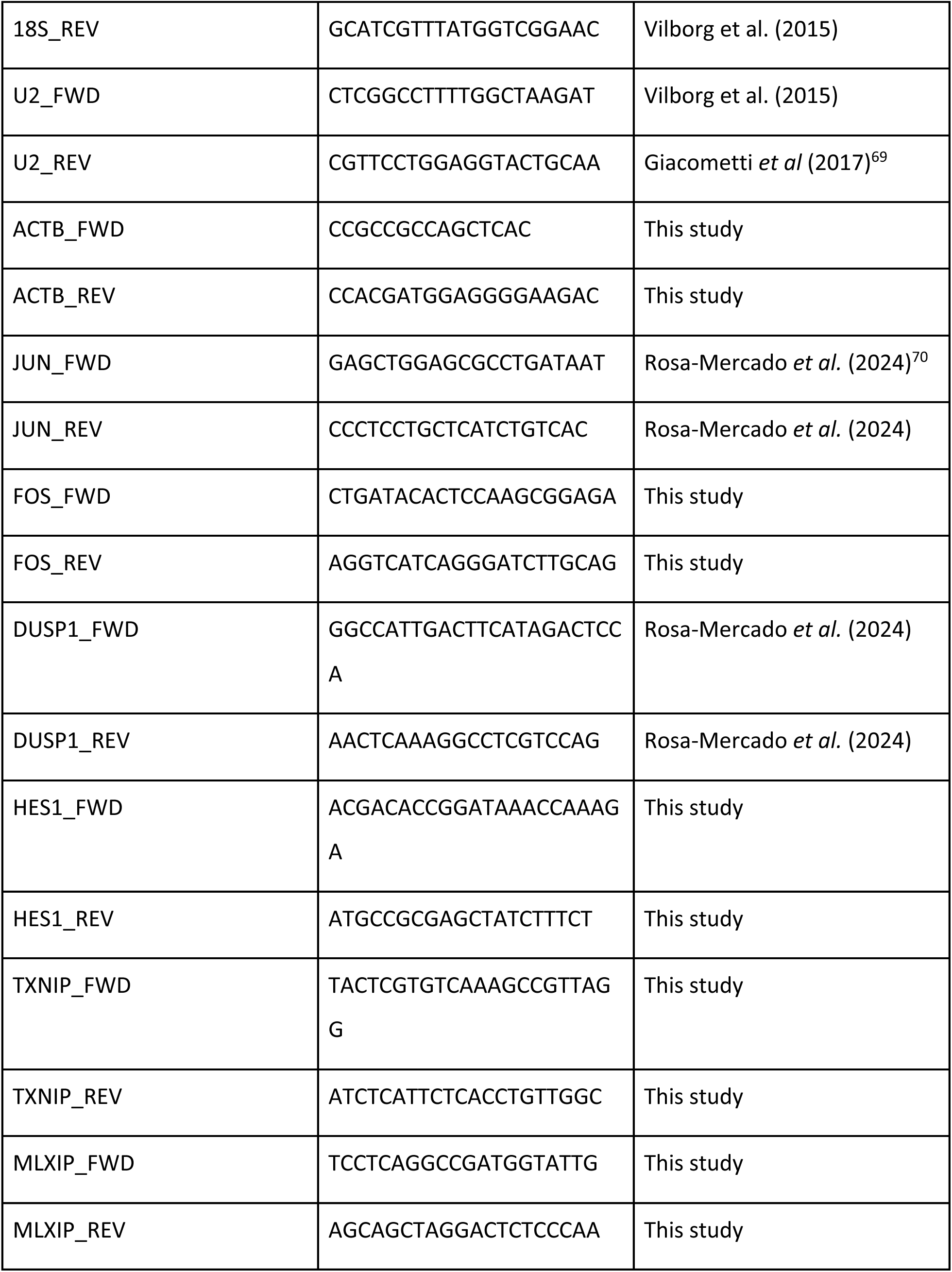
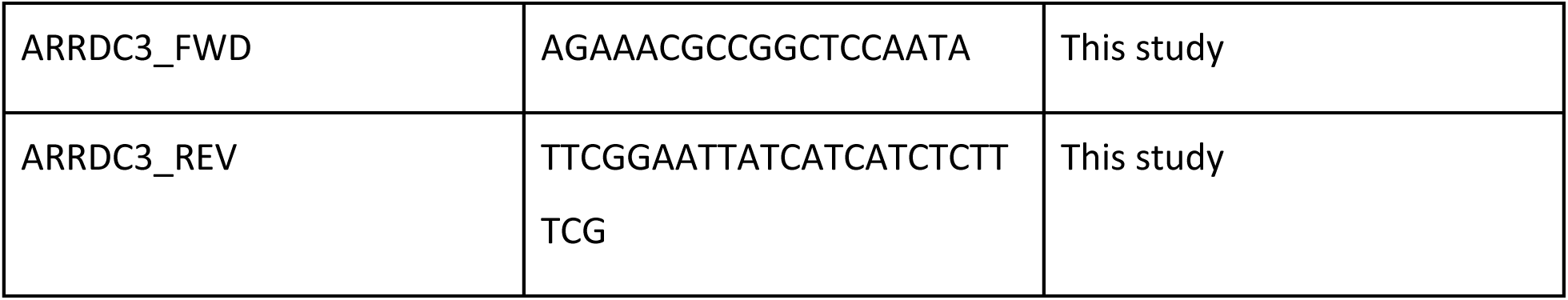

All experiments shown are representative of at least two biological replicates.

### Cell lines and maintenance

HEK293T cells (ATCC #CRL-3216) were acquired from ATCC. ZAK knockout HEK293T cells were generated in a previous study^19^. FreeStyle 293-F suspension cells (Thermo Fisher #R79007) were obtained from Thermo Fisher. Prior to use in experiments, all cell lines were passaged at least twice after thawing and were routinely checked for mycoplasma contamination. Adherent cells were cultured in DMEM (high glucose, L-glutamine, phenol red, sodium pyruvate - Thermo Fisher #11995073) supplemented with 10% FBS (Thermo Fisher #26140079). Cells were grown at 37 °C in a 5% CO_2_ humidified incubator and routinely passaged every 2-3 days. Suspension cells were cultured in HyCell TransFx-H medium (Cytiva, #SH30939) supplemented with 0.01% Poloxamer 188 (Sigma Aldrich, #P5556), 1x penicillin–streptomycin (Gibco, #15140122) and 3x GlutaMAX (Gibco, #35050061) at 37 °C, 5% CO_2_ and 80% humidity in a Multitron Cell (Infors HT) incubator.

### Cell seeding and drug treatment

Unless otherwise stated, for immunoblotting and RT-qPCR experiments, cells were seeded at 0.85e6 cells per well in 6-well plates the day before harvest to achieve ∼70-80% confluence at the time of harvest. Each well encompassed a single condition in a given experiment. The following day, the media of each well was exchanged for fresh growth media 1 h before drug treatment. For experiments involving pre-treatment with ZAK inhibitor or ISRIB, these drugs were added during the media change. Unless otherwise noted, ZAK inhibitor (nilotinib - APexBIO #A8232) and ISRIB (Sigma #SML0843) were added to a final concentration of 1 µM and 0.2 µM, respectively. 10 mM nilotinib solution in DMSO was purchased from APExBIO and aliquoted prior to storage at -80°C. ISRIB was dissolved to a 5 mM stock concentration in DMSO and stored at -80C.

Anisomycin (ANS - Sigma #A9789) and didemnin B (DDB - MedChemExpress #HY-105055) were dissolved in DMSO to a stock concentration of 94.2 mM and 10 mM respectively and stored at -80°C. 100 mM emetine (EME - Millipore #324693) and 200 mM L-Histidinol (Hist - Sigma #H6647) were prepared fresh the day of treatment in water. Sodium arsenite solution was obtained from Sigma (#106277). Unless otherwise stated, the following final drug concentrations were used in all experiments: anisomycin (0.38 µM), emetine (1.8 µM), didemnin B (0.5 µM), histidinol (4 mM), arsenite (100 µM). All drug treatments were performed by adding drug directly to the well to the final concentration, gently swirling the plate to mix, and incubating at 37°C for the appropriate treatment time. For all “untreated” conditions, volumes of DMSO equivalent to that used in the drug treatment conditions were added when applicable to serve as a vehicle control.

For arsenite-chase experiments, the media of each plate was exchanged for fresh growth media prior to incubation at 37°C for 1 h before drug treatment. The indicated elongation inhibitor was then added, and the plate was gently swirled to evenly distribute the drug. Following the given drug treatment time, arsenite was directly added to the drug-containing media and allowed to incubate at 37°C for the indicated chase time. The cells were then immediately harvested as described for the given assay.

### siRNA transfection

For experiments involving siRNA knockdowns, ∼0.6e6 cells were seeded onto 100 mm plates the day before transfection. The next day, cells were transfected with 250 pmol siRNA (Horizon Discovery) in OptiMEM using 37.5 µL Lipofectamine RNAiMAX (Thermo Fisher #13778150) according to the manufacturer’s instructions. The following day, the media of all transfected plates was exchanged for fresh growth media prior to incubation for another 2 days. Two days after the media change, cells were used in experimentation (i.e. drug treatment, etc.) and harvested as described.

### Cell harvest for immunoblotting

After drug treatment, each well was harvested by aspirating media, quickly washing with 1 mL PBS pH 7.4 (Thermo Fisher #10010049), and lysing cells with 150 µL ice-cold lysis buffer (RIPA buffer (Thermo Fisher #89901) supplemented with 1X Halt Protease and Phosphatase Inhibitor Cocktail (Thermo Fisher #71636), 10 mM sodium phosphate dibasic (Sigma #71636), 10 mM β-Glycerophosphate (Sigma #G9422), 42.5 units/mL benzonase (Sigma #E1014), and 1 mM TCEP (GoldBio #TCEP2)). The plate was swirled to evenly apply lysis buffer across the entire well before cells were scraped using a cell scraper. Cells were then resuspended gently using a pipette before transferring into a tube on ice to complete lysis. Lysates were then clarified by centrifuging at 8000 xg for 7 min at 4°C, and the clarified supernatant was transferred into a new tube on ice. Lysate supernatants were either used immediately or flash frozen at LN_2_ before storage at -80°C.

### Immunoblotting

Protein concentrations of clarified lysates were determined using BCA assay (Thermo Fisher #23227) according to the manufacturer’s instructions. Concentration-normalized SDS-PAGE samples were then prepared by mixing samples with lysis buffer and 6X Laemmli loading buffer before boiling at 95°C for 3-5 min. Samples were loaded onto 4-20% 26-well Criterion TGX Precast Midi Protein Gels (Bio-rad #5671095) before conducting electrophoresis at 150 V for 1 h in 1X Tris/Glycine/SDS buffer (Bio-rad #1610732). All gels included Precision Plus Protein Dual Color Standard (Bio-rad #1610374) protein ladder to keep track of protein size. Gels were transferred onto PVDF membranes (Bio-rad #1704273) using a Trans-blot Turbo Transfer System (Bio-rad) according to the manufacturer’s instructions. Membranes were blocked using 5% non-fat milk (Santa Cruz #sc-2324) dissolved in TBST (TBS supplemented with 0.1% Tween-20) for at least 30 min before addition of primary antibody in 5% milk in TBST. Membranes were incubated rocking with primary antibody at 4°C overnight. The following day, membranes were washed at 25°C, rocking in TBST for 5 min, four times total. Secondary antibody in 5% milk in TBST was then added for 1 h, rocking at 25°C for 1 h. Membranes were then washed at 25°C, rocking in TBST for 5 min, four times total. Membranes were incubated using SuperSignal™ West Pico PLUS (Thermo Fisher #34580) and Femto Maximum Sensitivity (Thermo Fisher #34095) Chemiluminescent Substrate prior to imaging in a ChemiDoc Imaging System (Bio-rad). All primary and secondary antibodies were used at concentrations as listed in the key resources table or at the manufacturer’s recommendation.

### Phostag immunoblotting

For phostag immunoblot analysis, concentration-normalized SDS-PAGE samples were prepared as described above. Samples were loaded onto 7% resolving acrylamide gels supplemented with 10.7 µM Phostag Acrylamide Reagent (Wako Fujifilm #AAL-107) and 21.3 µM MnCl_2_. EDTA-free pre-stained protein ladder (APExBio #F4005) was also loaded to track protein size. Electrophoresis was performed in 1X Tris/Glycine/SDS buffer (Bio-rad #1610732) at 100 V for 30 min and then 125 V for 2 h. Gels were then washed twice with 1X transfer buffer (25 mM Tris, 192 mM glycine, 10% methanol) supplemented with 1 mM EDTA (Thermo Fisher #AM9260G) to chelate residual Mn^2+^. Gels were then washed twice with 1X transfer buffer without EDTA. Gels were then transferred onto PVDF membranes (Bio-rad #1620177) using a standard wet-transfer protocol at 35 V at room temperature, overnight. Membranes were then blocked for at least 1.5 h in 5% milk in TBST before incubation with primary antibody at 4°C overnight. Membrane washing, secondary antibody incubation, and imaging were performed as described above.

### Cell harvest for ribosome pelleting

After drug treatment, each well was harvested by aspirating media, quickly washing with 1 mL PBS pH 7.4 (Thermo Fisher #10010049), and lysing cells with 150 µL ice-cold gradient lysis buffer (50 mM HEPES pH 7.4, 100 mM KOAc, 5% glycerol, 0.5% Triton X-100 (Sigma #T9284), 15 mM Mg(OAc)_2_, 1X Halt Protease and Phosphatase Inhibitor Cocktail (Thermo Fisher), 80 units TURBO DNase (Thermo Fisher #AM2239), 1 mM TCEP (GoldBio)). The plate was swirled to evenly apply lysis buffer across the entire well before cells were scraped using a cell scraper. Cells were then resuspended gently using a pipette before transferring into a tube on ice to complete lysis. Lysates were then clarified by centrifuging at 8000 xg for 7 min at 4°C, and the clarified supernatant was transferred into a new tube on ice. Lysate supernatants were either used immediately or flash frozen at LN_2_ before storage at -80°C.

RNA concentrations of lysates were determined using the Qubit RNA HS assay kit (Thermo Fisher #Q32852). Lysates were then concentration-normalized to the lowest concentration sample by mixing with gradient lysis buffer. 100 µL of normalized lysate was carefully layered on top of 100 µL 15% sucrose buffer/cushion (25 mM HEPES pH 7.4, 100 mM KOAc, 5 mM Mg(OAc)_2_, 15% sucrose, and 1 mM TCEP) before centrifuging in a MLA-130 fixed angle rotor (Beckman Coulter) at 300,000 xg for 22 min. Following centrifugation, the 200 µL supernatant was collected from each tube and mixed with 40 µL 6X Laemmli loading buffer for later SDS-PAGE analysis. 200 µL of pre-heated (95°C) nuclease-free water (Thermo Fisher) was then added to the tube to dissolve the pellet. After resuspension, this pellet fraction volume was transferred to a fresh tube and mixed with 40 µL Laemmli loading buffer. Corresponding input SDS-PAGE samples were generated by mixing equal volumes of normalized lysate and gradient lysis buffer before addition of 6X Laemmli loading buffer. All samples were heated at 95°C for 3-5 min before loading equal volumes of each fraction for SDS-PAGE and immunoblotting as described above.

### RNA extraction

After drug treatment, cells were harvested for RNA extraction by aspirating media and directly adding 1 mL TRIzol (Thermo Fisher #15596026) to each well. Cellular RNAs were then extracted following the TRIzol manufacturer’s protocol. In brief, cells were homogenized by gently pipetting up-and-down before transferring to a fresh tube. TRIzol lysates were incubated for 5 min prior to addition of chloroform and incubation for 2-3 min. The TRIzol-chloroform mixture was centrifuged at 12000 xg for 15 min at 4°C. The aqueous upper phase was transferred into a fresh tube containing Glycoblue coprecipitant (Thermo Fisher #AM9516). Isopropanol was added to the tube prior to inversion and incubation for 10 min at 4°C to precipitate RNAs. RNAs were pelleted by centrifugation at 12000 xg for 10 min at 4°C. The RNA pellet was then washed with 75% ethanol before centrifugation at 20000 xg for 5 min. The washed RNA pellet was then air-dried for 5-10 min before resuspension in nuclease-free water (Thermo Fisher #AM9932). Residual DNA was removed with TURBO DNase treatment at 37°C for 30 min (Thermo Fisher #AM2239). DNase was removed using a phenol:chloroform:isoamyl alcohol (Thermo Fisher #AM9732) extraction followed by an ethanol precipitation and wash. RNAs were resuspended in nuclease-free water and stored at -80°C. Concentrations were determined using a Nanodrop 8000 (Thermo Fisher).

### RT-qPCR

Reverse transcription was performed using the iScript cDNA synthesis kit (Bio-rad #1708891BUN) according to the manufacturer’s instructions. Resulting cDNAs were then diluted 1:2 in nuclease-free water (1 part cDNA, 2 parts water). qPCR reactions using diluted cDNAs were prepared using iTaq Universal SYBR Green Supermix (Bio-rad #1725121) according to the manufacturer’s instructions. Primers were used at a final concentration of 225 nM per reaction. qPCR was conducted using a QuantStudio 6 Flex real-time PCR system (Thermo Fisher) in a 384-well format. RT-qPCR data were analysed using a custom R script implementing the ddCt method using the geometric mean for 18S and U2 as a control^70,71^. RT-qPCR plots were plotted using Prism 10 (GraphPad).

### RNA-sequencing

RNA from drug-treated cells (in 6-well plates) was TRIzol-extracted as described above. Concentrations of TRIzol-extracted RNAs were measured by Nanodrop (Thermo Fisher). 10 µg of RNA was digested with 2 units Turbo DNase (Thermo Fisher) at 37°C for 30 min. DNase was removed by mixing reactions with an equal volume of RNAclean XP beads (Beckman Coulter #A63987) and washing with 80% ethanol two times prior to elution in nuclease-free water (Thermo Fisher). Eluted RNAs were used in RNA-sequencing library generation using the SMARTer Stranded Total RNA - HI Mammalian Sample Prep Kit (Takara #634882) according to the manufacturer’s instructions. Libraries were then pooled before 150 bp paired-end sequencing at Azenta.

### Data analysis

RNA-sequencing data was demultiplexed by Azenta. Adaptor sequences AGATCGGAAGAGCACACGTCTGAACTCCAGTCA and AGATCGGAAGAGCGTCGTGTAGGGAAAGAGTGT were trimmed from reads using cutadapt version 4.9^72^. Read pairs with at least one read shorter than 20 nt were discarded using cutadapt. The human reference genome GRCh38.p14 (release 46) and its associated annotation file were downloaded from Gencode. Genome index generation and read alignment was performed using STAR version 2.7.11b^73^ using the following arguments: “--runThreadN 36 --runMode alignReads --genomeDir $genome_index_path --readFilesIn $star_input_arg --readFilesCommand zcat -- outFilterMultimapNmax 1 --outFilterType BySJout --outSAMattributes All --outSAMtype BAM SortedByCoordinate --outFileNamePrefix $output_filename --quantMode GeneCounts”. Samtools version 1.20^74^ was used to generate, sort, and index .bam files. Read counts for each gene were generated using featureCounts version v2.0.6^75^. Downstream analysis was performed using R version 4.4.1^76^. Differential expression analysis of transcripts with more than 100 read counts was conducted using DESeq2 version 1.44.0^77^, with statistically significant genes selected based on a Benjamini-Hochberg adjusted p-value (padj) < 0.01. All DESeq2-calculated adjusted p-values were padded with an arbitrarily small number using .Machine$double.xmin in R (2.225074e-308) to allow proper plotting of differentially-expressed genes assigned a p-value of zero (due to their high significance). GSEA was performed in R using ClusterProfiler version 4.20.0^78–80^, using the human MSigDB Hallmark Gene Sets Collection^81,82^ imported by the R package msigdbr version 26.1.1^83^. Following DESeq2 analysis for each condition, processed lists of genes sorted by log2FoldChange were generated and inputted into ClusterProfiler’s GSEA function. Plots were generated using ggplot2 version 3.5.1^84^. Area-proportional Venn diagrams were generated using the DeepVenn webtool^85^.

### Polysome profiling and TCA precipitation

For non-siRNA transfection experiments, cells were seeded at 3.5e6 cells per plate on 100 mm plates the day before harvest to achieve ∼70-80% confluence at the time of harvest. For siRNA transfection experiments, cells were seeded and transfected as described above. On the day of harvest, the media was exchanged for fresh growth media 1 h before drug treatment, and then cells were treated with the appropriate inhibitor(s) dictated by the given experiment as described above.

After drug treatment, each plate was harvested by aspirating media, quickly washing with 5 mL PBS pH 7.4 (Thermo Fisher #10010049), and lysing cells with 200 µL ice-cold gradient lysis buffer (50 mM HEPES pH 7.4, 100 mM KOAc, 5% glycerol, 0.5% Triton X-100 (Sigma #T9284), 15 mM Mg(OAc)_2_, 1X Halt Protease and Phosphatase Inhibitor Cocktail (Thermo Fisher), 80 units TURBO DNase (Thermo Fisher #AM2239), 1 mM TCEP (GoldBio)). The plate was swirled to evenly apply lysis buffer across the entire well before cells were scraped using a cell scraper. Cells were then resuspended gently using a pipette before transferring into a tube on ice to complete lysis. Lysates were then clarified by centrifuging at 8000 xg for 7 min at 4°C, and the clarified supernatant was transferred into a new tube on ice. Lysate supernatants were either used immediately or flash frozen at LN_2_ before storage at -80°C.

A 10X gradient buffer solution (250 mM HEPES pH 7.5, 1 M KOAc, 50 mM Mg(OAc)_2_) was prepared, sterile filtered, and stored at room temperature. On the day of the experiment or on the day prior, 10-50% sucrose gradients were prepared by the following method. 10% sucrose and 50% sucrose solutions were prepared (1X gradient buffer, sucrose, 1 mM TCEP). 6 mL of 10% sucrose solution was added to 14 x 89 mm tubes (Seton Scientific #7030). A cannula was then used to add 6 mL 50% sucrose solution to the bottom of the tube, layering the 10% sucrose solution on top of the 50% sucrose solution. 10-50% sucrose gradients were then formed using a Biocomp Gradient Master prior to storage at 4C until use.

RNA concentrations of lysates were determined using the Qubit RNA HS assay kit (Thermo Fisher #Q32852). Lysates were then concentration-normalized to the lowest concentration sample by mixing with gradient lysis buffer. 300 µL of concentration-normalized lysates (40-60 µg total RNA) were layered on top of 10-50% sucrose gradients prior to centrifugation in a SW-41 swinging bucket rotor (Beckman Coulter #331362) at 40,000 rpm (274,400 xg) for 1 h 45 min at 4°C. Gradients were fractionated using a Bio-comp Piston Gradient Fractionator according to the manufacturer’s instructions. Trichloroacetic acid (TCA) was added to each fraction to a final concentration of ∼10% prior to overnight incubation at -20°C. Samples were then centrifuged at 20,000 xg for 15 min at 4°C prior to washing with 15% TCA and centrifugation at 20,000 xg for 15 min at 4°C. TCA pellets were then washed with ice-cold acetone prior to centrifugation at 20,000 xg for 10 min at 4°C, a total of two times. Pellets were then dried for 10 min at room temperature prior to dissolution in 30 µL sample buffer (2X Laemmli loading buffer, 100 mM Tris-HCl pH 8.0). Pellets were heated at 65°C for 10 min followed by 95°C for 10 min to ensure maximal dissolution into sample buffer. The samples were then used for immunoblotting as described above.

### Lysate preparation

FreeStyle 293-F cells were grown to a density of 1.0 x 10^6^ cells/mL and treated with 100 µM EME for 15 min. Cells were collected by centrifugation at 250 *x g* for 5 min and washed once with 1x PBS supplemented with 100 µM EME. Cells were resuspended in Extraction Buffer (20 mM HEPES/KOH pH 7.5, 150 mM KOAc, 5 mM MgCl_2_, protease-inhibitor mix (homemade), 1 mM DTT, 100 µM EME, 0.05% octaethylene glycol monododecyl ether) and passed five times through a 26-gauge needle. The lysate was cleared by centrifugation at 15,000 *x g* for 15 min. The resulting supernatant was directly subjected to cryo-EM sample preparation.

### Cryo-EM sample preparation

Carbon support R3/3 300 mesh copper grids with a 3 nm amorphous carbon layer (Quantifoil) were charged by treatment with 1-pyrenemethylamine (PMA; Sigma Aldrich #401633) as described before^86^. In brief, grids were submerged in 50 mM PMA dissolved in DMSO for 30 s followed by two consecutive washes in isopropanol and ethanol for 5 s each. Grids were allowed to dry for 30 min. 3.5 µL lysate were applied to a grid, incubated for 45 s and blotted for 3 s before plunge freezing in liquid ethane using a Vitrobot (FEI) at 5 °C and 85% humidity.

### Cryo-EM data collection and processing

Cryo-EM data for the EME cell lysate was collected on a Titan Krios (FEI) at 300 kV equipped with a Falcon 4i direct election detector (Themo Fisher) and a Selectris X energy filter (Thermo Fisher) with 20 eV slit width. A nominal pixel size of 0.727 Å/pixel, a defocus range of -0.5 to -3.5 µm and a total dose of 40 e^-^/Å^2^ were used.

Movies were motion corrected using MotionCor2^87^ and contrast-transfer function (CTF) parameters were estimated using CTFFIND4^88^. Micrographs with an estimated resolution above 5 Å were removed. In total 36,242 micrographs remained (**Figure S8A**). For particle picking, a crYOLO (v1.7.6)^89^ model was newly trained based on manual picks from a random subset of 100 low-pass filtered micrographs (PhosaurusNet architecture, standard settings). 765,906 particles were picked and extracted in Relion (v4.0.1)^90^ 4x binned with 160 pixels box size. Particles were imported into cryoSPARC (v4.6.0)^91^ and subjected to 2D classification. Classes corresponding to 80S ribosomes were selected (**Figure S8B**) and an initial reconstruction was obtained after a homogeneous refinement using an ab initio generated reference. After 3D classification in cryoSPARC, good classes containing EME in a TI-4/-5 translocation intermediate state (107,490 paricles; 39%) were selected as representative for EME-stalled and potentially leading ribosomes (**Figure S8C**). For these particles the box was iteratively expanded including multiple steps of re-centering and refinement to a 5x binned box size of 840 pixels. After removal of particles close to the edge of the micrographs 89,851 particles (32%) remained. These particles were subjected to focused 3D classification in cryoSPARC using a wide spherical mask around density for a collided, neighboring ribosome. The class with the most robust density was selected (19,698 particles; 7%). These particles were re-extracted 5x binned with an expanded box size of 960 pixels and removal of out-of-bounds particles reduced their number to 18,231 (6.5%). Another focused 3D classification was performed in cryoSPARC using a spherical mask around the collided ribosome density and a class with the least orientation biased neighboring 80S density was selected (6,664 particles; 2.4%). Due to the relatively low number of particles and the arising orientation bias further box expansion and recentering on the neighboring ribosome was not possible. Instead, we chose another class from our initial unmasked 3D classification that corresponds to a pre-translocation state with partial eEF2 density and also density for a leading, stalled ribosome as representative for the collided ribosome (48,490 particles; 17%). Here, we also performed multiple steps of box expansion, re-centering and refinement up to a 6x binned box size of 1020 pixels. Once out-of-bound particles were removed 34,205 particles (12%) remained. These particles were subjected to focused 3D classification in cryoSPARC using a wide spherical mask around the mRNA entry site and a class with the best neighboring density was selected (12,064 particles; 4%). The box was again expanded including multiple steps of re-centering and refinement to a 6x binned box size of 1200 pixels. Removal of edge particles left 10,491 particles (4%). To improve the quality of the stalled ribosome density we performed particle subtraction using an inverse mask leaving only the stalled ribosome. Subsequently, we used a regular mask around the stalled ribosome for non-uniform refinement in cryoSPARC. To mitigate the eminent orientation bias we rebalanced particles based on orientation in cryoSPARC. This reduced the orientation bias but also the number of particles to 6,997 (2.5%). To reconstruct the EME induced disome collision we then used the sorted EME stalled ribosome (6,664 particles) and the sorted pre-translocation trailing ribosome (10,491 particles) to generate a low-resolution binned map where the focused non-uniform refined collided ribosome (6,997 particles) served as fitting aid for the EME stalled ribosome. To gain high resolution information that was lost due to the low number of final particles and the orientation bias – especially in the collided ribosome class – we selected the initial particles from the first unmasked 3D classification (39% EME stalled; 17% pre-translocation collided) and extracted them unbinned at 640 pixels box size in Relion. We imported these particles into cryoSPARC and performed a homogeneous refinement with global CTF and per-particle defocus refinement. This resulted in overall resolutions of 2.55 and 2.92 Å for the EME TI-4/TI-5 and the pre-translocation states, respectively (**Figure S9**). Finally, the densities were local resolution filtered. These high-resolution densities were then backfitted into the low-resolution EME disome composite map to generate a high-resolution composite map (**Figure S8C**). The box was trimmed in Phenix (v2.0-5936)^92^ to produce a manageable file size. Lastly, to further improve the resolution of the EME binding site we performed particle subtraction on the EME TI-4/TI-5 particles using an inverse mask leaving only the 40S body followed by local refinement and local resolution filtering using a mask around the 40S body. This resulted in an overall resolution of 2.59 Å for the 40S body of the EME stalled ribosome (**Figure S9**).

### Model building

The models for the EME stalled TI-4/TI-5 state and the collided pre-translocation state were initially assembled individually. First, for the stalled ribosome PDB:8GLP^93^ was used as base. 60S, 40S head and body were rigid body docked individually. They were then manually adjusted. tRNAs and mRNA from PDB:8GLP were also docked and adjusted the same way. The model for eEF2 was taken from PDB:6Z6M^94^, fitted and manually adjusted. The initial model for the P-stalk was taken from PDB:9I2D^95^ and rigid body fitted and manually adjusted. Models for the P-stalk proteins uL10 and uL11 were taken from the AlphaFold 2 database^96^ rigid body docked and manually adjusted as much as the local resolution allowed for. The L1-stalk and ribosomal protein uL1 were taken from PDB:6Z6M and rigid body docked. Adjustment of the L1 stalk was limited by the local resolution as well. For EME, the monomer (ligand ID:34G) was manually fitted and adjusted.

For the collided ribosome, again PDB:8GLP was used as base for the ribosome with 60S, 40S head and body fitted individually and then adjusted manually. A/P and P/E tRNAs as well as the mRNA were taken from PDB:9RPV^19^, fitted and manually adjusted. The C-terminal domain of EDF1 was also isolated from PDB:9RPV, fitted and manually adjusted to the degree the local resolution allowed for. 18S rRNA helix h16 was also taken from PDB:9RPV, fitted and adjusted to match the conformation induced by EDF1.

Both ribosomes were first individually real-space refined against the composite map and adjusted before they were combined to produce the disome model which was then real-space refined again.

All manual adjustments were performed in Coot (v9.8.96)^97^ and all real-space refinements were done using Phenix (v2.0-5936)^92^ (Table S1).

## Data availability

Cryo-EM maps and molecular models were deposited at the Electron Microscopy Data Bank (EMDB) and the Protein Data Bank (PDB), respectively. They are available via the following accession codes: EMD-57348 and 29SQ (Emetine disome); EMD-57345 (Emetine stalled 80S - TI4/TI5 State); EMD-57346 (Collided 80S - pre-translocation State); EMD-57347 (Emetine stalled 80S -TI4/TI5 State - 40S body local refinement).

RNA-sequencing data is available on the NCBI GEO repository under the accession number GSE329747. Code used to analyze RNA-sequencing data is available on Zenodo (DOI: 10.5281/zenodo.19957590).

Uncropped immunoblot images have been deposited in Mendeley Data and are publicly accessible as of the date of publication at the following DOI: 10.17632/5t8swbpbxm.1

